# Hepatic mTORC1 signaling activates ATF4 as part of its metabolic response to feeding and insulin

**DOI:** 10.1101/2021.05.02.442369

**Authors:** Vanessa Byles, Yann Cormerais, Krystle Kalafut, Victor Barrera, James E. Hughes Hallett, Shannan Ho Sui, John M. Asara, Christopher M. Adams, Gerta Hoxhaj, Issam Ben-Sahra, Brendan D. Manning

## Abstract

**Objective:** The mechanistic target of rapamycin complex 1 (mTORC1) is dynamically regulated by fasting and feeding cycles in the liver to promote protein and lipid synthesis while suppressing autophagy. However, beyond these functions, the metabolic response of the liver to feeding and insulin signaling orchestrated by mTORC1 remains poorly defined. Here, we determine whether ATF4, a stress responsive transcription factor recently found to be independently regulated by mTORC1 signaling in proliferating cells, is responsive to hepatic mTORC1 signaling to alter hepatocyte metabolism.

**Methods:** ATF4 protein levels and expression of canonical gene targets were analyzed in the liver following fasting and physiological feeding in the presence or absence of the mTORC1 inhibitor rapamycin. Primary hepatocytes from wild-type or liver-specific *Atf4* knockout (*LAtf4^KO^*) mice were used to characterize the effects of insulin-stimulated mTORC1-ATF4 function on hepatocyte gene expression and metabolism. Both unbiased steady-state metabolomics and stable-isotope tracing methods were employed to define mTORC1 and ATF4-dependent metabolic changes. RNA-sequencing was used to determine global changes in feeding-induced transcripts in the livers of wild-type versus *LAtf4^KO^* mice.

**Results:** We demonstrate that ATF4 and its metabolic gene targets are stimulated by mTORC1 signaling in the liver in response to feeding and in a hepatocyte-intrinsic manner by insulin. While we demonstrate that *de novo* purine and pyrimidine synthesis is stimulated by insulin through mTORC1 signaling in primary hepatocytes, this regulation was independent of ATF4. Metabolomics and metabolite tracing studies revealed that insulin-mTORC1-ATF4 signaling stimulates pathways of non-essential amino acid synthesis in primary hepatocytes, including those of alanine, aspartate, methionine, and cysteine, but not serine.

**Conclusion:** The results demonstrate that ATF4 is a novel metabolic effector of mTORC1 in liver, extending the molecular consequences of feeding and insulin-induced mTORC1 signaling in this key metabolic tissue to the control of amino acid metabolism.

## 1. Introduction

The liver is a central effector of systemic metabolic flexibility with a critical role in coupling shifts in glucose, lipid and amino acid metabolism to nutrient fluctuations that occur with fasting and feeding [1]. At the molecular level, the mechanistic target of rapamycin complex 1 (mTORC1) is at the heart of a nutrient sensing network that integrates nutrient availability with hormonal signals to calibrate cellular metabolism [2]. Such signal integration by mTORC1 occurs through two convergent G-protein switches, the amino acid-sensing pathway regulating the Rag GTPases and hormonal regulation of the tuberous sclerosis complex (TSC) protein complex and Rheb GTPase [3]. In the liver, mTORC1 activity is suppressed with fasting and activated with feeding to coordinate the shift between energy-producing catabolic processes and energy-consuming anabolic processes accompanying these two states [4, 5]. Feeding-induced activation of hepatic mTORC1 stimulates protein and lipid synthesis while suppressing autophagy [4, 6–8]. Conversely, fasting diminishes mTORC1 activity in the liver, relieving its inhibitory signals on fatty acid oxidation, ketogenesis, and autophagy [5, 7]. However, our current understanding of the metabolic effectors and processes downstream of mTORC1 signaling in the liver is incomplete.

Through studies largely performed in cell culture models, the mTORC1 protein kinase complex has been established to exert acute metabolic control through phosphorylation events on a growing list of direct downstream substrates. These mTORC1 targets include the canonical targets S6K1 and 4EBP1 to promote protein synthesis [9] and ULK1 and TFEB to suppress autophagy and lysosome biogenesis [10], as well as the S6K1-specific target CAD to stimulate *de novo* pyrimidine synthesis [11, 12]. Additionally, mTORC1 promotes metabolic alterations by engaging a downstream transcriptional network of genes encoding key metabolic enzymes [3]. Over the last decade, mTORC1 signaling has been found to stimulate a coordinated transcriptional response through regulation of specific transcription factors, including HIF1α to promote glucose uptake and glycolysis [13–16], SREBP1 and 2 to stimulate lipogenesis [4, 6, 15, 17, 18], and NFE2L1/NRF1 to support proteasome synthesis [19]. The regulation of these transcription factors downstream of mTORC1 has largely been characterized in proliferating cells in response to growth factors or oncogenic signaling. Aside from SREBP1c activation [4, 6, 20], the transcriptional effectors contributing to the metabolic response orchestrated by mTORC1 in the liver and terminally differentiated hepatocytes in response to feeding and insulin are poorly defined.

Activating transcription factor 4 (ATF4) is the best-characterized downstream effector of the integrated stress response (ISR), which is coordinated by four stress-sensing kinases - GCN2, PERK, HRI, and PKR - that converge to phosphorylate the translation initiation factor eIF2α at serine 51 [21]. Phosphorylation of eIF2α results in global attenuation of cap-dependent translation along with selective translation of ATF4, mediated by altered regulation of translation of upstream open reading frames (uORFs) in the 5’ untranslated region (UTR) of the ATF4 transcript [21]. Elevated levels of ATF4 directly stimulate the expression of genes involved in adaptation to the cellular stresses initiating the ISR [22]. Beyond its canonical role in the ISR, ATF4 has also been found to be activated by anabolic signals, including insulin, downstream of mTORC1 in cell-based studies [23–26]. Pro-growth signals that activate mTORC1 stimulate ATF4 translation independently of the ISR to induce a subset of its gene targets, thereby promoting specific mTORC1-stimulated metabolic processes, including the synthesis of protein, purine nucleotides, and glutathione [24, 26]. However, our knowledge of ATF4 function in the liver is limited, and it is unknown whether physiological activation of mTORC1 promotes ATF4 function to alter hepatocyte metabolism.

In the current study, we find that physiological activation of mTORC1 by feeding in the liver and insulin in primary hepatocytes stimulates the activation of ATF4 and increased expression of ATF4-dependent gene targets. We find that despite robust mTORC1-ATF4-mediated regulation of previously established anabolic targets involved in serine synthesis and one carbon metabolism in response to insulin that ATF4 is dispensable for mTORC1-stimulated *de novo* nucleotide synthesis in primary hepatocytes. Furthermore, our study reveals that cultured primary hepatocytes synthesize little if any serine, even under serine/glycine deprivation conditions. Unbiased metabolite profiling and subsequent stable isotope-tracing experiments revealed that insulin-mTORC1-ATF4 signaling induces the synthesis of S-adenosylmethionine (SAM) in the methionine cycle and cystathionine in the trans-sulfuration pathway in primary hepatocytes. In addition, we find that insulin signaling through mTORC1 and ATF4 induces synthesis of the non-essential amino acids aspartate and alanine, albeit without detectable changes to expression of the specific transaminases involved. RNA-seq analyses of livers from control (*Atf4^fl/fl^*) and liver-specific *Atf4* knockout (*LAtf4^KO^*) mice revealed that a subset of ATF4 target genes, largely related to amino acid metabolism, were controlled by ATF4 in response to feeding. Thus, our study demonstrates that ATF4 is a novel downstream effector of physiological mTORC1 activation in the liver that contributes to the broader anabolic cellular program downstream of insulin and mTORC1 signaling.

## 2. Materials and Methods

### 2.1. Mice and Diets

All mice were maintained at the Harvard T.H. Chan School of Public Health and procedures were performed with prior approval and in accordance with the guidelines set forth by the Harvard Institutional Animal Care and Use Committee. Wild-type C57/BL6J mice (males age 6-8w) were purchased from Jackson Laboratories. *Atf4^fl/fl^* mice were generated as previously described by the insertion of loxP sites flanking exons 2 and 3 and backcrossed to a C57/BL6J background for 9 generations [27]. *Atf4^fl/fl^* mice were crossed with C57/BL6J mice expressing the Albumin-Cre transgene (Jackson Labs) to generate liver-specific ATF4 knockout mice (*LAtf4^KO^*), as previously described [28]. For fasting/feeding studies, mice were fasted for 12h during the light cycle and either euthanized or fed a high carbohydrate diet (Harlan Teklad, Basal Mix adjusted for fat, TD.88122) for 6-12h in the dark cycle, as previously described[29]. Vehicle (5% Tween-80, 5% PEG-400, DMSO in 1x PBS) or 10 mg/kg rapamycin (LC laboratories) was injected i.p. 30 minutes prior to feeding. For all studies, mice were anesthetized with isoflurane, and when necessary, blood was collected retro-orbitally in EDTA-coated microtubes for plasma isolation. Animals were humanely euthanized, and organs were snap-frozen in liquid nitrogen, with the left lobe of the liver used for all assays. For tunicamycin treatment, mice were either injected i.p. with vehicle (150mM dextrose in 1x PBS) or tunicamycin at 1mg/kg for 6h.

For glucose tolerance tests, *Atf4^fl/fl^* and *LAtf4^KO^* male mice were fasted for 16h overnight (7p.m.-11a.m) and injected with glucose at 1mg/kg. For insulin tolerance tests, mice were fasted for 6h during the day (8a.m.-2p.m.) and injected with insulin 0.75U/kg (Eli Lily HumulinR) dissolved in PBS plus protease free BSA. Blood glucose was monitored over time using the OneTouch® Ultra glucometer.

### 2.2. Primary Hepatocyte Isolation and Culture

Primary wild-type hepatocytes were isolated from C57/BL6J male and female mice at 7-10 weeks of age (Jackson Labs). Portal vein perfusion of buffer A (10mM HEPES, 150mM NaCl, 5mM KCl, 5mM glucose, and 2.5mM sodium bicarbonate, 0.5mM EDTA, pH 8.0) was performed for 5-7 minutes at a rate of 5ml/minute followed by buffer B (buffer A minus EDTA, with 35mM CaCl_2_ and Liberase TM, Sigma-Aldrich, 5401127001) for 5-7 minutes at a rate of 5ml/minute. Livers were placed in Dulbecco’s Modified Eagle Medium (DMEM 4.5g/L glucose, w/o sodium pyruvate; Corning, 15-017-CV), 2.5% FBS, Pen/Strep, and Glutamax (ThermoFisher, 35050061) or Glutagro (Corning, 25-015-CI), and the liver capsule was disrupted to release hepatocytes. Hepatocytes were centrifuged at 1000rpm for 5 minutes and then resuspended in 10ml DMEM, 9ml Percoll (Sigma-Aldrich, P4937), and 1ml 10x PBS. Following a spin at 1000rpm for 7 minutes, hepatocytes were washed once with media, spun at 1000rpm for 5 minutes, and then resuspended in 10ml media per mouse. Hepatocytes from 3-5 mice were pooled and viable hepatocytes, determined by Trypan blue exclusion and cell counts, were plated at 1.25-1.5×10^6^ cells/well in 6-well collagen-coated dishes (BioCoat, Corning) or 2.5×10^6^ cells/6cm dishes for metabolite profiling or tracing experiments (BioCoat, Corning). After 4-6h, media was changed to serum-free media (DMEM, 2mM L-glutamine) overnight followed by insulin stimulation (100nM human insulin, Sigma-Aldrich, I9278), with 30-minute vehicle, rapamycin, or Torin1 pre-treatment, where indicated. For siRNA delivery, hepatocytes were transfected with 25-125nM siRNA using Lipofectamine RNAiMax (5μl 6-well, 10μl 6cm) 3h after plating. After 4h, the media was changed to serum-free media overnight or to low serum media (DMEM, 1% FBS, 2mM L-glutamine). For siRNA experiments extending beyond 24h (*Eif4ebp1* and *Eif4ebp2*), hepatocytes were maintained in 1nM dexamethasone with 1% FBS until incubation in serum-free media. For adenoviral delivery, hepatocytes were infected with Ad5-CMV-eGFP (U. Iowa Vector Core) or mouse Ad5-CMV-ATF4-eGFP (Vector Biolabs, ADV-253208) 4h after plating at an MOI of 10. After overnight incubation (∼16h), the media was changed to low serum media (1% FBS) and incubated for another 8h (total= ∼24h).

Primary human hepatocytes were obtained from Lonza (Cat#HUCPG, Lot#HUM4252) and were thawed according to the manufacturer’s instructions into William’s E Medium (without phenol red, ThermoFisher), 5% FBS, Pen/Strep, 15mM HEPES, and 100nM dexamethasone and plated in 24-well dishes for RNA (375,000/well) and 12-well dishes for protein (750,000). After 6h of plating, hepatocytes were serum starved in William’s E Medium, 10nM dexamethasone overnight. Media was changed the following morning to serum free William’s E Medium (without dexamethasone) for treatment with inhibitors and 100nM insulin for 6h.

### 2.3. Cell Lines

Hepa1-6 and AML-12 murine hepatocyte cell lines were obtained from the ATCC and maintained in DMEM, 4.5g/L glucose without sodium pyruvate (Thermo Scientific) and supplemented with Pen/Strep and Glutagro (Corning) and DMEM-F12 1:1, (ThermoFisher Scientific) supplemented with Pen/Strep, Glutagro (Corning), insulin-transferrin-selenium (ThermoFisher Scientific) and dexamethasone (100nM, Sigma Aldrich), respectively.

### 2.4. Reagents

After dissolving in DMSO, tunicamycin (Sigma-Aldrich, T7765) was used at 2μg/ml, rapamycin (EMD Millipore, 553210) at 20nM, and Torin1 (Tocris, 4247) at 500-750nM. Control non-targeting pool (D-001810-10-50) and ONTarget Plus SMARTpool siRNAs against mouse *Atf4* (L-042737-01-0020, 25nM), *Mthfd2* (L-042690-01-0020, 25nM), *Eif4ebp1* (L-058681-01-005, 125nM), and *Eif4ebp2* (L044972-01-005, 125nM) were purchased from Horizon Discovery/Dharmacon. Lipofectamine RNAiMAX was purchased from ThermoFisher Scientific. ^15^N-glutamine-amide (490024) and ^15^N-glutamine-amine (486809) were purchased from Sigma-Aldrich. 3-^13^C-serine (CLM-1572), ^13^C_5_-methionine (CLM-893-H), and U-^13^C_6_-glucose (CLM-1396) were purchased from Cambridge Isotope Laboratories. High glucose DMEM without cystine and methionine (21013024) was purchased from ThermoFisher Scientific. DMEM without serine and glycine (U.S. Biologicals, D-9802-01) was dissolved in water and supplemented with 4.5g/L glucose, sodium bicarbonate, and phenol red followed by filter sterilization.

### 2.5. Immunoblotting

Protein extracts were prepared from tissues and cells using RIPA buffer (50mM Tris-Cl pH 7.4, 150mM NaCl, 1% IGEPAL, 0.5% sodium deoxycholic acid, 0.1% SDS, 1mM EDTA, 10mM NaF, 10mM sodium pyrophosphate, 1mM β-glycerophosphate, and 1mM sodium orthovanadate, Sigma protease inhibitor P8340). Extracts from liver tissue were prepared by homogenizing liver pieces (∼25-50mg) in RIPA buffer with protease inhibitor, Halt Phosphatase Inhibitor Cocktail (ThermoFisher Scientific, 78420), and Phosphatase Inhibitor Cocktail I (Sigma-Aldrich) using the Red Lysis Kit (Next Advance) and Next Advance Bullet Blender (speed 8, for 3 minutes). Protein concentrations were determined using a BCA assay kit (Thermo Scientific) or a detergent compatible Bradford assay (ThermoFisher Scientific). Equal amounts of protein (15-20µg) were separated by SDS-PAGE, transferred to nitrocellulose membranes, and immunoblotted with indicated primary antibodies. Primary antibodies: ATF4 (BioLegend, 693902 used in primary hepatocytes and liver tissue), MTHFD2 (Proteintech,12270-1-AP), MTHFD2 (Abcam, ab151447), PHGDH (Sigma-Aldrich, HPA021241), PSAT1 (Proteintech, 20180-1-AP), PSPH (Proteintech, 14513-1-AP), ATF4 (Proteintech, 10835-1-AP, used in liver tissue), GOT2 (Proteintech, 14800-1-AP), SREBP-1 (Santa Cruz, sc-13551), β-actin (Sigma-Aldrich A5316), α-Tubulin (Sigma-Aldrich, T6074), phospho (P)-S6K1 T389 (Cell Signaling Technologies (CST), 9234 used in cells), P-S6K1 T389 (CST, 97596 used in liver tissue) S6K1 (CST, 2708), ATF4 (CST, #11815 used in cell lines), PERK (CST, 3192), P-eIF2α S51 (CST, 3597), eIF2α (CST, 9722), P-AKT S473 (CST, 4060), Pan-AKT (CST, 4691), P-S6 S240/44 (CST, 2215), S6 (CST, 2217), 4E-BP1 (CST, 9644), 4E-BP2 (2845), P-CAD S1859 (CST, 12662), CAD (CST, 11933), P53 (CST, 32532), CHOP (CST, 5554), GAPDH (CST, 5174) and GOT1 (CST, 34423). Secondary antibodies: anti-rabbit IgG, HRP-linked (CST, 7074), anti-mouse IgG, HRP-linked (CST 7076), anti-rat IgG, HRP-linked (CST, 7077), IRDye 800CW donkey anti-mouse IgG (H+L) (LI-COR, 926-32212) and donkey ant-rabbit IgG (H+L) (LI-COR, 925-32213). Immunoblots were developed by ECL (West Pico or Femto, Thermo Scientific) or with a LI-COR Odyssey CLx imaging system (LI-COR Biosciences). For ATF4 measurement in livers, 40-50ug of protein was separated on 4-15% TGX Criterion gels and transferred to nitrocellulose membranes. After transfer, membranes were rinsed with 1x TBS (without Tween-20) and blocked with 5% milk in 1x TBS (without Tween-20) for 1h at room temperature. Membranes were then incubated at 4°C overnight with ATF4 antibody (BioLegend) at 1:1000. After washing with 1x TBST, anti-rat HRP secondary antibody (CST) was used at 1:2000. A list of commercially available antibodies tested in livers and cell lines is included in Supplementary Table 1.

### 2.6. Gene Expression Analysis

RNA was isolated from cells (6-well dishes) or liver pieces (∼10-15mg) using the RNeasy Plus Mini kit (Qiagen). For livers, samples were homogenized in RLT buffer and suspended in an equal volume of 50% ethanol prior to application to the RNeasy columns. RNA (0.5-1µg) was reverse transcribed using the Advanced cDNA Synthesis Kit (Bio-Rad). Skirted plates and iTaq SYBR green for qPCR were purchased from Bio-Rad. QPCR analysis was performed in biological duplicates or triplicates and with duplicate technical replicates. Analysis was performed using the CFX96 Real Time PCR Detection System. Samples were normalized to Rplp0 (36b4) for ^ΔΔ^Ct analysis using the Bio-Rad CFX96 software. Primer sequences are listed in Supplementary Table 2.

### 2.7. Steady State Metabolite Profiling and Targeted Metabolic Flux Analysis

Primary mouse hepatocytes from 3-5 mice were pooled and plated at 2.5×10^6^ cells/6cm dish in triplicate or quadruplicate. Hepatocytes were washed twice with media lacking the amino acid used as the tracer (Gln, Ser, or Met) and incubated in the same media containing tracer for the last 30 min (^15^N-glutamine-amide (2mM)) or 1h (^15^N-glutamine-amine (2mM), 3-^13^C_1_-serine (400μM), or ^13^C_5_-methionine (200μM)) of insulin stimulation . Metabolites were extracted with 80% methanol (HPLC grade) for 15 minutes at -80°C, with cells subsequently scraped off of plates into 80% methanol on dry ice and placed into 15ml conical tubes. After 5-minute centrifugation at maximum speed, supernatants were transferred to 50ml conical tubes. A second extraction was performed on the remaining pellet with 500μl of ice-cold 80% methanol, centrifuged, and pooled with the first extraction in 50ml conical tubes. After the final extraction, remaining insoluble pellets were resuspended in 500μl of 8M urea in 10mM Tris-Cl pH 8.0 and shaken at 60°C for 1h. Protein concentrations were measured for sample normalization using a BCA or Bradford assay. Metabolite extracts were dried under a stream of nitrogen gas using an N-EVAP (Organomotion Associates, Inc.).

For metabolite tracing into the nucleotides of total cellular RNA, primary hepatocytes plated in 6-well dishes were labeled with ^15^N-glutamine-amide (2mM) for 6h. RNA was isolated using the RNeasy Plus Mini Kit (Qiagen) according to the manufacturer’s instructions. RNA (4-5μg) was heated to 100°C for 3 minutes and rapidly cooled in an ice-water bath. Samples were brought up to 50μl and subsequently digested with 1μl of Nuclease P1 in 50mM sodium acetate buffer (New England Biolabs, M0660) for 2h at 45°C. Samples were then neutralized with 1/10 volume of 1M ammonium bicarbonate (made fresh) and digested with 2μl of 0.25U/μl Phosphodiesterase I from *Crotalus adamanteus* venom (Sigma-Aldrich, P3243) for 2h at 37°C. Samples were then extracted in 80% methanol and dried under nitrogen gas using an N-EVAP.

Dried-down metabolites were re-suspended in 30μL of HPLC-grade water and 5μL of sample was injected for liquid chromatography/mass spectrometry (LC/MS) using a 6500 QTRAP hybrid triple quadrupole mass spectrometer (AB/SCIEX) coupled to a Prominence UFLC HPLC system (Shimadzu) with Amide XBridge HILIC chromatography (Waters) via selected reaction monitoring (SRM, Supplementary Table 3) and polarity switching between positive and negative modes [30]. For steady state profiling, selected reaction monitoring of a total of 254 polar metabolites was analyzed [30]. Parameters for software analysis were as previously described [11]. Peak areas from the total ion current for each metabolite SRM transition were integrated using MultiQuant v2.0 software (AB/SCIEX). Peak areas were normalized to protein levels from insoluble pellets obtained during the metabolite extraction. As a control, an unlabeled sample was run in parallel to account for natural abundance for comparison to m+1 labeled isotopologues. For RNA tracing experiments, the unlabeled signal was subtracted from the labeled isotopologues before calculating the fractional enrichment. For steady state metabolomic profiling, KEGG enrichment analysis was performed using MetaboAnalyst software and the heatmap of significantly altered metabolites was generated using Morpheus software (Broad Institute of MIT and Harvard).

### 2.8. RNA-Sequencing

RNA was isolated from the livers of *Atf4^fl/fl^* and *LAtf4^KO^* mice (n=5/group) with the Qiagen RNeasy Mini kit. RNA-sequencing was performed at the Dana-Farber Cancer Institute Sequencing Core. Strand-specific libraries were generated with 500ng of RNA using TruSeq library preparation kit (Illumina, San Diego, CA). The cDNA libraries were multiplexed and sequenced using Illumina NextSeq 500 with single-end 75bp read length parameters. Adapter sequences were trimmed off sequencing adaptors and low-quality regions by using cutadapt [31]. Trimmed reads were aligned to UCSC build mm10 of the Mus musculus genome, using STAR [32]. After the counts were collected, differential gene expression analysis was performed using DE-Seq2, which calculated fold change and adjusted p-values [33]. Lists of differentially expressed genes were examined for gene ontology (GO) and KEGG term enrichment with clusterProfiler [34]. Gene set enrichment analysis was performed using MSigDB annotated gene sets (Broad Institute of MIT and Harvard). The complete RNA-seq data can be found at GEO under the accession number GSE173578.

### 2.9. Statistical Analysis

All statistical analyses were performed using GraphPad Prism 8.0 using either Student’s t-tests, One-way ANOVA (with Holm-Sidak post-hoc analysis) or Two-way ANOVA (with Holm-Sidak pos-hoc analysis), as indicated in each figure legend. *p<0.05, **p<0.01, ***p<0.001, ****p<0.0001.

### 2.10. Immunoblot quantification and schematic generation

Western blots were quantified using either LICOR or ImageJ software. Schematic figures were generated using BioRender software.

## 3. Results

### 3.1. Physiological stimulation of mTORC1 signaling activates ATF4 and expression of established gene targets

To determine if mTORC1 activation in the liver stimulates an increase in ATF4 protein with physiological feeding, mice were subjected to a 12h light-cycle fast followed by 6h refeeding in the dark cycle with a high carbohydrate diet. As expected, refeeding potently stimulated mTORC1 activity as assessed by phosphorylation of its downstream targets S6K1 and 4E-BP1, the latter reflected in an upward electrophoretic mobility shift (**Figure 1A**). Additionally, we observed a feeding-induced, rapamycin-sensitive increase in phosphorylation of the S6K1 target CAD, the first enzyme of de novo pyrimidine synthesis [11, 12], as well as the namesake S6K1 substrate ribosomal protein S6. Importantly, feeding-induced mTORC1 signaling also stimulated increased protein expression of ATF4, which could be blunted by administration of rapamycin prior to feeding. Refeeding mice 12h in the dark cycle also led to induction of liver ATF4 protein levels in an mTORC1-dependent manner with overall reductions in mouse to mouse variability (Figure S1A). Importantly, unlike ATF4, eIF2α phosphorylation in the liver, indicative of ISR induction, was not consistently affected by fasting, feeding, or rapamycin, supporting previous studies in cell-based models that mTORC1 can activate ATF4 occurs independent of the ISR [24, 26]. As we had found previously that mTORC1 activation stimulates expression of genes encoding enzymes of the serine synthesis pathway (*Phgdh, Psat, Psph*) and mitochondrial one-carbon metabolism (*Mthfd2*) through ATF4 [24], we assessed whether these genes were influenced by feeding and mTORC1 signaling in the liver. Indeed, along with *Atf4* transcripts, feeding induced expression of these genes in a rapamycin-sensitive manner (**Figure 1B**). However, rapamycin did not affect the ability of feeding to suppress expression of the gluconeogenic gene *Pepck*, indicative of a lack of global effects from this treatment on the feeding response.

**Figure 1.**
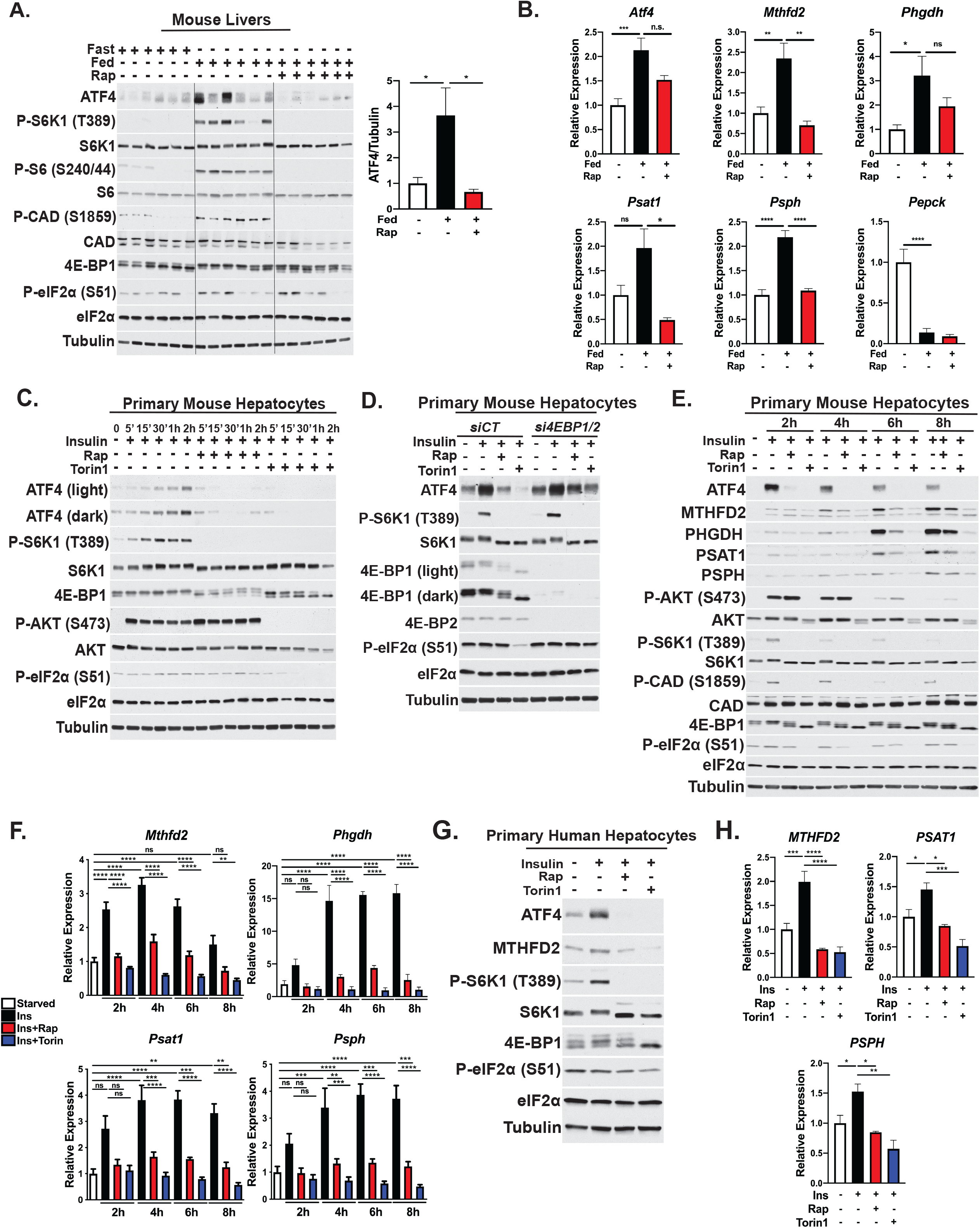
Feeding and insulin induce hepatic ATF4 and established gene targets via mTORC1 signaling. **(A,B)** Eight-week old male mice were fasted for 12h and refed a high carbohydrate diet for 6h following pretreatment with vehicle or 10mg/kg rapamycin (n=6/group). **(A)** Liver lysates were immunoblotted to assess mTORC1 signaling and ATF4, with mean ATF4 protein to tubulin quantification plotted ± SEM and normalized to the fasted group (right). **(B)** Liver gene expression plotted as mean ± SEM relative to the fasted group. **(C)** Immunoblot analysis of primary mouse hepatocytes serum starved overnight and treated with 100nM insulin following a 30-minute pretreatment with vehicle (DMSO), 20nM rapamycin, or 750nM Torin1 for the indicated time points. **(D)** Immunoblot analysis of primary mouse hepatocytes transfected with control (siCT) or a combination of *Eif4ebp1 (4ebp1)* and *Eif4ebp2* (*4ebp2*) siRNAs, followed by overnight serum starvation and treatment for 8h with 100nM insulin following a 30-minute pretreatment with vehicle, 20nM rapamycin or 500nm Torin1. **(E)** Immunoblot analysis of primary mouse hepatocytes treated for the indicated time points as in (C). **(F)** Gene expression in cells treated as in (E) plotted as mean ± SEM relative to serum-starved cells (n=4 independent experiments). **(G)** Immunoblot analysis of primary human hepatocytes serum starved overnight and subsequently treated with 100nM insulin following a 30-minute pretreatment with vehicle (DMSO), 20nM rapamycin, or 500nM Torin1 for the indicated time points. **(H)** Gene expression in cells treated as in (G) plotted as mean ± SEM (n=3) relative to the serum-starved cells. *p<0.05, **p<0.01, ***p<0.001, ****p<0.0001 (one-way ANOVA).

Given that the liver contains both parenchymal and non-parenchymal cells, we sought to characterize the cell-intrinsic regulation of ATF4 and its transcriptional targets by mTORC1 in primary mouse hepatocytes. Over a time course of insulin stimulation, where Akt is fully activated within 5 min and mTORC1 signaling by 30 min, and increase in ATF4 protein levels was evident by 30 min, peaking at 2 hours (**Figure 1C**). Insulin-stimulated mTORC1 signaling and induction of ATF4 were both blocked by rapamycin or the kinase domain inhibitor of mTOR Torin1, which has more potent suppressive effects on mTORC1 targets and global protein synthesis [35]. Additionally, while there was no observable change in eIF2α phosphorylation with insulin or rapamycin treatment, we did observe that Torin1 abrogated the basal eIF2α phosphorylation signal with time (**Figure 1C**), suggesting that the stronger effects of Torin1 on ATF4 and its gene targets observed (see below) might be due to combined effects on both mTORC1 signaling and the ISR. For this reason, in all subsequent experiments, we use rapamycin instead of, or in parallel to, Torin1 treatment, to focus on mTORC1-specific effects on ATF4 regulation and function.

We next sought to determine how mTORC1 signaling stimulates hepatic ATF4 activation in response to insulin. A previous report demonstrated that mTORC1-dependent regulation of ATF4 occurs through phosphorylation and inactivation of the translational repressors 4E-BP1 and 4E-BP2 [25, 36]. Indeed, silencing of 4E-BP1 and 4E-BP2 in primary hepatocytes with siRNAs resulted in elevated basal ATF4 protein levels and resistance to rapamycin and Torin1 treatment relative to hepatocytes transfected with control siRNAs (**Figure 1D**). Interestingly, while there was no effect of rapamycin on eIF2α phosphorylation under either condition, knockdown of 4E-BP1/2 rescued the effects of Torin1 treatment on basal eIF2α phosphorylation, indicating that these effects are likely linked to the 4EBP1/2-dependent attenuation of protein synthesis reported with Torin1 treatment in other settings [35], which could increase amino acid availability and minimize protein load on the ER, thus decreasing basal activation of the ISR.

To further characterize the hepatocyte-intrinsic effects of the mTORC1-ATF4 axis, we performed insulin time course experiments analyzing the transcript and protein levels of the aforementioned ATF4 targets. Based on time course experiments, the peak of ATF4 protein production upon insulin stimulation was at 2h, with levels steadily declining over the time course up to 8h (**Figure 1C, E**). Induction of *Mthfd2*, *Phgdh*, *Psat1*, and *Psph* transcript levels was detected by 2h insulin stimulation, peaking at 4h, while corresponding protein levels were robustly induced by 6h (**Figure 1E, F**, with protein quantified in Figure S1B). Importantly, both mRNA and protein levels of these gene targets were sensitive to rapamycin and Torin1, with the latter demonstrating more potent effects accompanying decreased eIF2α phosphorylation. In addition to primary hepatocytes, the commonly used murine hepatocyte cell lines, Hepa1-6 and AML-12, also displayed insulin-stimulated increases in ATF4 and expression of these gene targets that was blocked by mTOR inhibitors (Figure S1C-D). Finally, in primary human hepatocytes, insulin induced ATF4 protein in an mTORC1-dependent manner, without effects on eIF2α phosphorylation (**Figure 1G**), and this regulation correlated with stimulated expression of *MTHFD2*, *PSAT1*, and *PSPH* transcripts (**Figure 1H**). Together, these findings define ATF4 as a novel downstream target of physiological mTORC1 activation with feeding and insulin in the liver and cell-autonomously in hepatocytes.

### 3.2. ATF4 is required for insulin-mTORC1 signaling to induce the expression of serine synthesis and one carbon metabolism enzymes in hepatocytes

Both the basal and insulin-stimulated expression of MTHFD2, PHGDH, PSAT1, and PSPH transcripts and proteins were abrogated in primary hepatocytes with siRNA-mediated knockdown of *Atf4* (**Figure 2A,B**). As an orthogonal approach, we generated mice with liver-specific deletion of Atf4 (*LAtf4^KO^*), using previously described *Atf4^fl/fl^* mice crossed to mice expressing the Cre recombinase from the liver-specific albumin promoter [27, 28]. In primary hepatocytes isolated from control *Atf4^fl/fl^* mice, insulin induced MTHFD2, PHGDH, PSAT1, and PSPH protein and transcripts in a rapamycin-sensitive manner, but this regulation was greatly reduced in the *LAtf4^KO^* hepatocytes (**Figure 2C,D**). Of note, the *LAtf4^KO^* hepatocytes, like those with siRNA knockdown of *Atf4*, displayed enhanced insulin-stimulated mTORC1 signaling, perhaps indicative of loss of ATF4-dependent targets that negatively regulate mTORC1 [37–40]. To confirm the specificity of the ATF4-dependent responses, primary hepatocytes from *LAtf4^KO^* mice were infected with adenoviruses expressing either GFP-encoding control (AdGFP) or an ATF4 cDNA (AdATF4). The markedly diminished expression of *Mthfd2*, *Phgdh*, *Psat1*, and *Psph* transcripts in the *LAtf4^KO^* hepatocytes was fully restored with AdATF4. Furthermore, ectopic expression of ATF4 in wild-type primary hepatocytes resulted in substantial overexpression of ATF4 that correlated with enhanced MTHFD2 and PSAT1 protein levels (**Figure 2G**). As the AdATF4 adenovirus lacks the endogenous 5’UTR of ATF4, which is required for its regulation by mTORC1 [24–26], the enhanced expression of ATF4, MTHFD2 and PSAT1 induced by AdATF4 was resistant to rapamycin treatment. Together, these results establish that the mTORC1-ATF4 axis regulates these gene targets in primary hepatocytes with potential implications for metabolic control of serine and purine nucleotide synthesis, as observed in proliferative settings [24].

**Figure 2.**
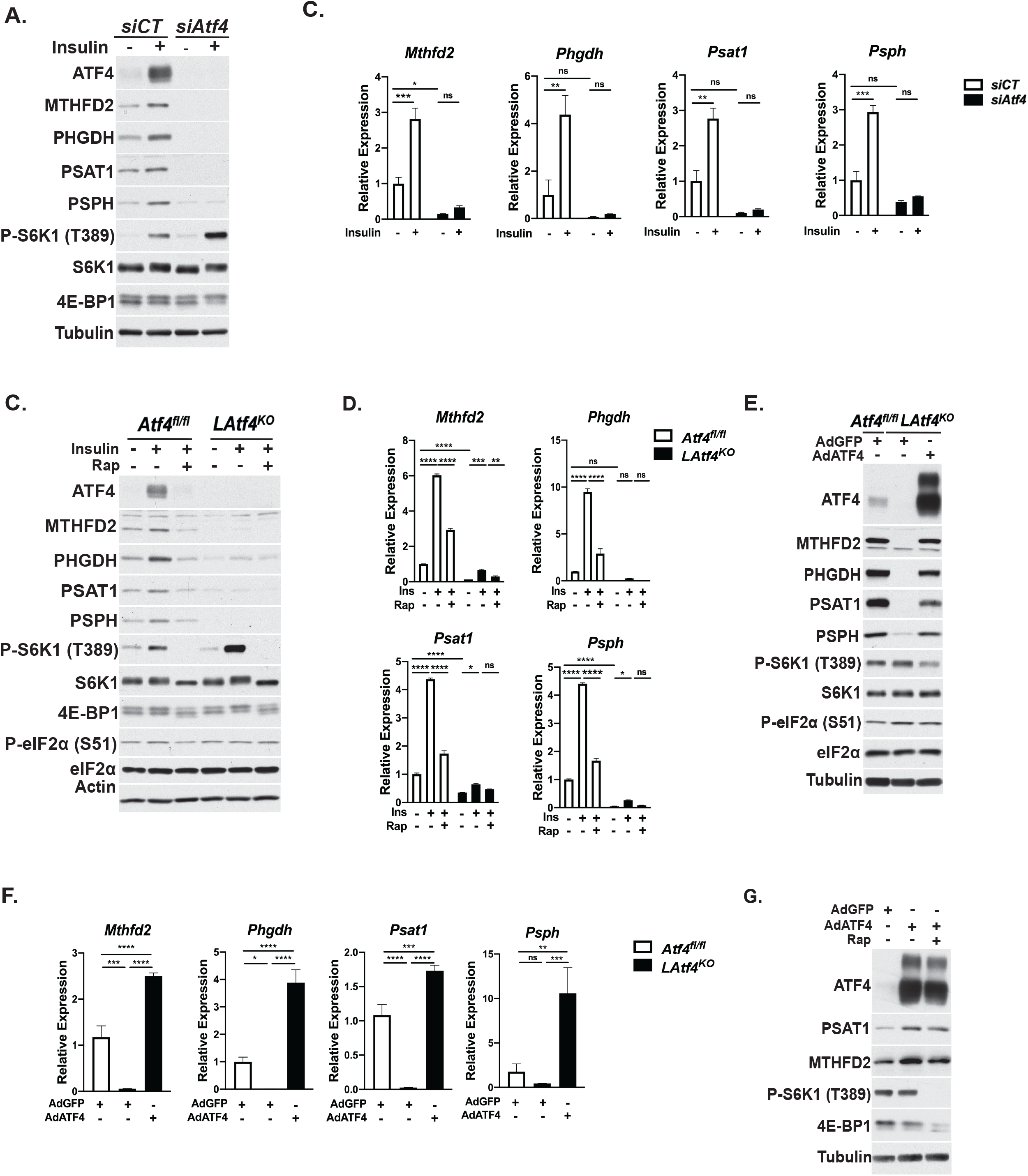
ATF4 is both necessary and sufficient for insulin-mTORC1 signaling to induce select gene targets. **(A)** Immunoblot analysis of primary mouse hepatocytes transfected with control (siCT) or *Atf4*-targetting siRNAs followed by overnight serum starvation and treatment with 100nM insulin for 8h. **(B)** Gene expression in cells treated as in (A) plotted as mean ± SEM relative to the *siCT*-transfected serum-starved cells (n=3 independent experiments). **(C)** Immunoblot analysis of *Atf4^fl/fl^* and *LAtf4^KO^* primary mouse hepatocytes serum starved overnight and subsequently treated with 100nM insulin for 6h following a 30-minute pretreatment with vehicle (DMSO) or 20nM rapamycin. **(D)** Gene expression in cells treated as in (C) plotted as mean± SEM relative to the *Atf4^fl/fl^* serum-starved cells (n=3). **(E)** Immunoblot analysis of *Atf4^fl/fl^* and *LAtf4^KO^* primary mouse hepatocytes infected with AdGFP or AdATF4 (MOI=10) for 24h. **(F)** Gene expression from a representative experiment of cells treated as in (E) plotted as mean± SEM relative to the *Atf4^fl/fl^* AdGFP sample (n=3). **(G)** Immunoblot analysis of WT primary mouse hepatocytes infected with AdGFP or AdATF4 (MOI=10) for 24h followed by treatment with vehicle (DMSO) or 20nM rapamycin for 8h. *p<0.05, **p<0.01, ***p<0.001, ****p<0.0001 (two-way ANOVA (B,D) or one-way ANOVA (F)).

### 3.3. mTORC1 stimulates hepatocyte nucleotide synthesis in an ATF4-independent manner

In proliferating cells, mTORC1 regulates *de novo* purine nucleotide synthesis, in part, through ATF4-dependent regulation of MTHFD2 and the mitochondrial tetrahydrofolate cycle, which supplies one carbon formyl units derived from the carbon 3 atom of serine to the purine ring [24] (**Figure 3A**). While primary hepatocytes are terminally differentiated cells with minimal DNA synthesis, they still have a high demand for nucleotides to produce rRNA needed for ribosome biogenesis in this secretory organ. Indeed, we found that insulin stimulated an mTORC1-dependent increased flux through the *de novo* purine synthesis pathway in primary hepatocytes, as measured by ^15^N-glutamine-amide tracing into the newly synthesized free pool of cellular purine nucleotides, including IMP (m+2), ATP (m+2) and GTP (m+3), the latter of which acquires a third ^15^N atom from the glutamine amide (**Figure 3B**). Likewise, 3-^13^C_1_-serine tracing, which specifically measures incorporation of the two formyl units into the purine ring, also revealed that insulin stimulates increased *de novo* purine synthesis in an mTORC1-dependent manner (**Figure S2A**). We next determined the effects of insulin-mTORC1 signaling on flux through *de novo* purine synthesis into the total RNA pool of primary mouse hepatocytes, of which approximately 80% represents rRNA [41]. The fractional enrichment for RNA-derived purines labeled with ^15^N-glutamine-amide via de novo synthesis during an 8h stimulation with insulin was surprisingly low (<4%). However, an insulin-stimulated, mTORC1-dependent increase in the fractional enrichment of *de novo* synthesized AMP (m+2) and GMP (m+3) within RNA was observed (**Figure 3C**). Unexpectedly, siRNA-mediated knockdown of *Atf4* in primary hepatocytes did not impair the insulin-stimulated increase in ^15^N-glutamine-amide or 3-^13^C_1_-serine flux into free pools of purine nucleotides (**Figure 3D** **and Figure S2B**). Furthermore, direct silencing of *Mthfd2* also failed to impact the ability of insulin to stimulate de novo purine synthesis (**Figure S2B**). Likewise, *Atf4* knockdown had only minor impact on the fractional enrichment of *de novo* synthesized AMP (m+2) and GMP (m+3) derived from total hepatocyte RNA (**Figure 3E**).

**Figure 3.**
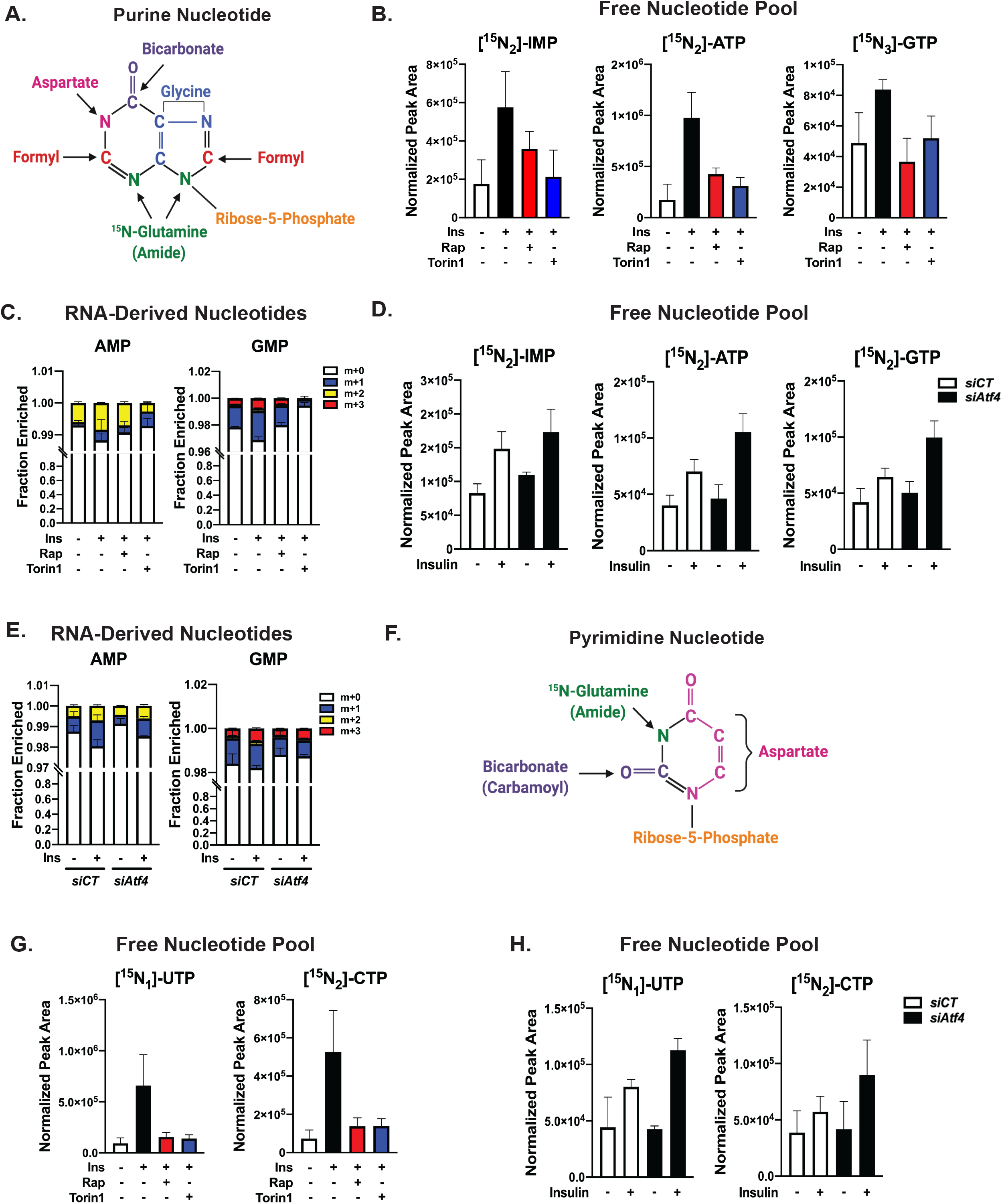
Insulin-mTORC1 signaling induces hepatic *de novo* nucleotide synthesis in an ATF4-independent manner. **(A)** Schematic of purine nucleotide highlighting carbon and nitrogen sources for *de novo* synthesis. **(B)** Protein normalized peak areas of ^15^N-glutamine-amide tracing (2mM, last 30 minutes) into free labeled purine nucleotide pools from primary mouse hepatocytes serum starved overnight and subsequently treated with 100nM insulin following a 30-minute pretreatment with vehicle (DMSO), 20nM rapamycin, or 750nM Torin1 for 8h. Data are plotted as mean ± SD and are representative of two independent experiments performed in quadruplicate. **(C)** Fractional enrichment of ^15^N-glutamine-amide tracing into RNA-derived purine nucleotide isotopologues in primary mouse hepatocytes serum starved overnight and subsequently treated with 10nM insulin following a 30 minute pretreatment with vehicle (DMSO), 20nM rapamycin, or 500nM Torin1 for 8h in medium containing 2mM ^15^N-glutamine-amide tracer. Data are plotted as mean ± SD and are representative of two independent experiments performed in triplicate. **(D)** Protein normalized peak areas of ^15^N-glutamine-amide tracing (2mM, last 30 minutes) into free labeled purine nucleotide pools from primary mouse hepatocytes transfected with control (siCT) or *Atf4*-targetting siRNAs 4h after plating, followed by overnight serum starvation and overnight treatment with 10nM insulin. Data are plotted as mean ± SD and are representative of two independent experiments performed in quadruplicate. **(E)** Fractional enrichment of ^15^N-glutamine-amide tracing into RNA-derived purine nucleotide isotopologues in cells transfected as in (D), followed by overnight serum starvation and treatment with 10nM insulin in medium containing 2mM ^15^N-glutamine-amide tracer for 6h. Data are plotted as mean ± SD and is representative of three independent experiments performed in triplicate. **(F)** Schematic of pyrimidine nucleotide highlighting carbon and nitrogen sources for *de novo* synthesis. **(G)** ^15^N-glutamine-amide tracing (2mM, last 30 minutes) into free labeled pyrimidine nucleotide pools in primary mouse hepatocytes treated and plotted as in (B). **(H)** Protein normalized peak areas of ^15^N-glutamine-amide tracing (2mM, last 30 minutes) into free labeled pyrimidine nucleotide pools from primary mouse hepatocytes transfected with control (siCT) or *Atf4*-targetting siRNAs 4h after plating, followed by overnight serum starvation and overnight treatment with 100nM insulin. Data are plotted as mean ± SD and are representative of two independent experiments performed in quadruplicate.

The effects of insulin-mTORC1 signaling on *de novo* synthesis of pyrimidine nucleotides was also assessed in primary hepatocytes. Consistent with the stimulated phosphorylation and activation of CAD by S6K1 downstream of mTORC1 detected in liver and primary hepatocytes (**Figure 1A,E**) [11, 12], insulin stimulated ^15^N-glutamine-amide flux into free pools of UTP (m+1) and CTP (m+2) and RNA-derived UMP (m+1) and CMP (m+2) in an mTORC1-dependent manner (**Figure 3F,G** **and Figure S2C**). Based on the role of ATF4 in regulating non-essential amino acid metabolism in other cellular settings [22, 25, 26, 42], we assessed de novo pyrimidine synthesis upon *Atf4* knockdown. Rather than inhibiting insulin-stimulated pyrimidine synthesis, loss of ATF4 moderately enhanced ^15^N-glutamine-amide flux into the free pools of pyrimidines and had no observable effects on the fractional enrichment of labeling into RNA-derived pyrimidines (**Figure 3H** **and Figure S2D**). Thus, insulin-mTORC1 signaling induces de novo synthesis of both purine and pyrimidine nucleotides in primary hepatocytes through mechanisms independent of ATF4.

### 3.4. Lack of evidence for regulated serine synthesis in primary hepatocytes

The *de novo* serine synthesis pathway (SSP) is a metabolic branchpoint in glycolysis comprised of three successive steps catalyzed by the PHGDH, PSAT1, and PSPH enzymes, with the product serine contributing to multiple biosynthetic processes, in addition to protein and nucleotide synthesis (**Figure 4A**). To measure SSP activity in primary hepatocytes, we used ^15^N-glutamine-amine tracing to capture the transamination of 3-phosphohydroxypyruvate to 3-phosphoserine catalyzed by PSAT1 and serine generation following dephosphorylation by PSPH (**Figure 4A**). Despite robust regulation of PHGDH, PSAT1, and PSPH expression by mTORC1-ATF4 signaling, described above, insulin failed to stimulate ^15^N-glutamine-amine flux into the free pool of labeled serine (m+1) in primary hepatocytes (**Figure 4B**). One possibility is that the newly synthesized serine was being rapidly utilized for protein synthesis, however, treatment with the protein synthesis inhibitor cycloheximide during the 1h labeling did not result in accumulation of labeled serine. Surprisingly, this 1h labeling yielded a fractional enrichment of labeled serine of less than 5% in primary hepatocytes, which was unaltered by insulin, mTORC1 inhibitors, or cycloheximide (**Figure 4C**). This result was unchanged with labeling times from 30 min up to 6h, indicating that flux through the SSP is unusually low in primary hepatocytes (**Figure S3A,B**), even under these conditions where the expression of SSP enzymes are induced by insulin through mTORC1 and ATF4. Consistent with this disconnect between pathway enzyme regulation and SSP flux, siRNA-mediated knockdown of *Atf4* had no effect on the relative abundance or fractional enrichment of labeled serine (m+1) in primary hepatocytes (**Figure 4D,E**).

**Figure 4.**
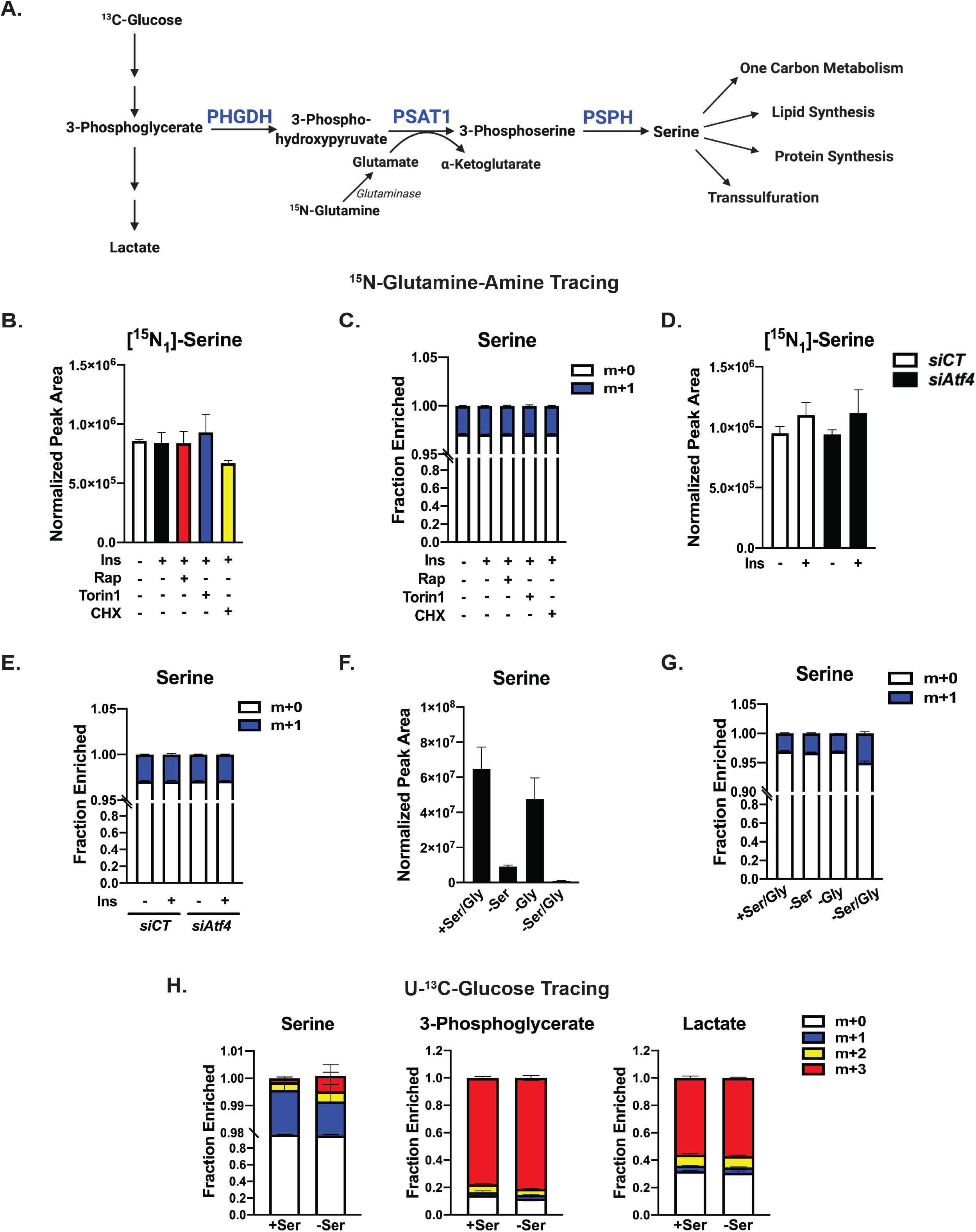
Lack of evidence for regulated serine synthesis in hepatocytes. **(A)** Schematic of the serine synthesis pathway. **(B)** Protein normalized peak areas of ^15^N-glutamine-amine (2mM, last 1h) tracing into labeled serine in primary mouse hepatocytes serum starved overnight and subsequently treated with 100nM insulin following a 30-minute pretreatment with vehicle (DMSO), 20nM rapamycin, or 750nM Torin1 for 8h, or where indicated, cycloheximide (50µM) was added with tracer for the last 1h. Data are plotted as mean ± SD and are representative of two independent experiments performed in quadruplicate. **(C)** Fractional enrichment of serine isotopologues from the experiment in (B). **(D)** Protein normalized peak areas of ^15^N-glutamine-amine (2mM, last 1h) tracing into labeled serine in primary mouse hepatocytes transfected with control (siCT) or *Atf4*-targetting siRNAs, followed by overnight serum starvation and overnight treatment with 10nM insulin. Data are plotted as mean ± SD and are representative of two independent experiments performed in quadruplicate. **(E)** Fractional enrichment of serine isotopologues from the experiment in (D), plotted as mean ± SD. **(F)** Protein normalized peak areas of intracellular serine from primary mouse hepatocytes following incubation in serine and/or glycine-free medium plus 2.5% dialyzed FBS for 24h. Data are plotted as mean ± SD and are representative of two independent experiments performed in triplicate. **(G)** Fractional enrichment of serine isotopologues in primary mouse hepatocytes treated as in (F) and labeled with ^15^N-glutamine-amine (2mM, last 1h). Data are plotted as mean ± SD and is representative of two independent experiments performed in triplicate. **(H)** Fractional enrichment of isotopologues from primary mouse hepatocytes following incubation in serine-rich or serine-free medium + 2.5% dialyzed FBS for 20h with 10mM ^13^C_6_-glucose. Data plotted as mean ± SD and is representative of two independent experiments performed in quadruplicate.

We next determined whether primary hepatocytes were capable of activating SSP flux in response to serine and glycine deprivation, as other cell types do [43–45]. 24-h serine starvation of primary hepatocytes resulted in a marked reduction in intracellular serine levels that was further reduced with combined serine and glycine starvation (**Figure 4F**). However, this serine depletion had little effect on the fractional enrichment of newly synthesized serine (m+1) detected with ^15^N-glutamine-amine labeling (**Figure 4G**). As glucose is the primary source of carbon for serine synthesis, we also utilized U-^13^C-glucose tracing into serine. Despite robust (80%) labeling of 3-phosphoglycerate (m+3), the glycolytic intermediate precursor to the SSP, the fractional enrichment of glycolysis-derived labeled serine (m+3) was only slightly increased with serine deprivation and still accounted for less than 1% of total cellular serine (**Figure 4H**). This is in contrast to another product downstream of 3-phosphoglycerate, lactate, which was nearly 60% m+3 labeled over the same duration. Together, these results indicate that primary hepatocytes in culture synthesize very little serine, even when the enzymes of the SSP pathway are elevated or cells are deprived of exogenous serine.

### 3.5. ATF4 regulates methionine metabolism in hepatocytes

As the above studies on nucleotide and serine synthesis revealed that hepatocytes are distinct from other cellular systems in the control of these processes, we employed unbiased steady state metabolomics to identify potential insulin-stimulated metabolic changes dependent on ATF4. This analysis revealed that several metabolites were significantly elevated (p<0.05) upon insulin treatment in control hepatocytes but not those with siRNA-mediated knockdown of *Atf4* (**Figure 5A**, with the top 6 metabolites shown graphically in **Figure 5B**, **Supplementary Table 4**). The metabolites induced most strongly with insulin in an ATF4-dependent manner included those related to the transsulfuration pathway and the methionine cycle, including cystathionine and S-adenosylmethionine (SAM) (**Figure 5A-C**). Indeed, enrichment analysis of the insulin-ATF4 regulated metabolites revealed an overrepresentation of KEGG metabolite sets for amino acid metabolism, including that of cysteine and methionine (**Figure S5A**). Additionally, we observed decreased levels of reduced glutathione upon ATF4 knockdown, with modestly increased levels of oxidized glutathione disulfide (GSSG) (**Figure 5A, Figure S5B**). We recently demonstrated that the mTORC1-ATF4 axis stimulates the production of glutathione through induction of SLC7A11 expression and increased cystine uptake in proliferating cells [26]. However, *Slc7a11* expression was not regulated by insulin or ATF4 in primary hepatocytes, nor were the glutathione synthesis enzymes *Gclc* and *Gclm* (**Figure S4C**).

**Figure 5.**
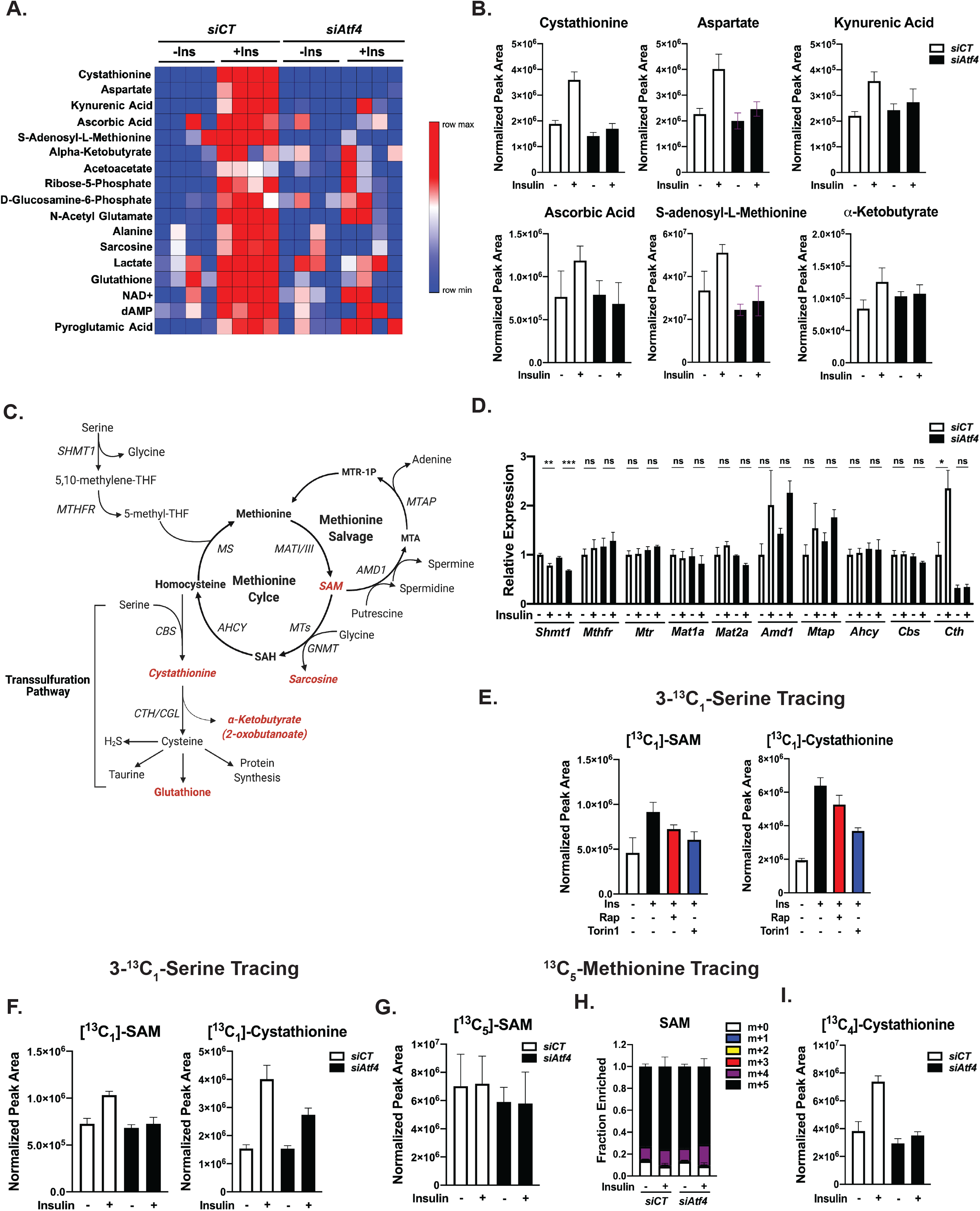
Insulin stimulates metabolic flux into the methionine cycle and transsulfuration pathway through ATF4. **(A)** Steady state metabolomic profiling from primary mouse hepatocytes transfected with control (siCT) or *Atf4*-targetting siRNAs followed by 100nM insulin stimulation for 6h performed in quadruplicate. The heat map displays a rank order of metabolites significantly (p<0.05) induced with insulin in the control cells but not those with *Atf4* knockdown. **(B)** Protein normalized peak areas of the top 6 metabolites (A) plotted as mean ± SD. **(C)** Schematic of the interconnections between serine and tetrahydrofolate metabolism, the methionine cycle and the transsulfuration pathway. Metabolites found to be significantly induced by insulin in an ATF4-dependent manner in (A) are highlighted in red. **(D)** Gene expression analysis of primary mouse hepatocytes transfected with control (siCT) or *Atf4*-targetting siRNAs followed by overnight serum starvation and treatment with 100nM insulin for 8h. Data are plotted as mean ± SEM relative to *siCT*-transfected serum starved cells (n=3 independent experiments). **(E)** Protein normalized peak areas of 3-^13^C-serine (400µM, last 1h) tracing into labeled SAM and cystathionine in primary mouse hepatocytes serum starved overnight and subsequently treated with 100nM insulin following a 30-minute pretreatment with vehicle (DMSO), 20nM rapamycin, or 500nM Torin1 for 8h. Data are plotted as mean ± SD and are representative of two independent experiments performed in triplicate. **(F)** Protein normalized peak areas of 3-^13^C-serine tracing (400µM, last 1h) in primary mouse hepatocytes transfected with control (siCT) or *Atf4*-targetting siRNAs followed by overnight serum starvation and treatment with 100nM insulin for 8h. Data are plotted as mean ± SD and are representative of two independent experiments performed in triplicate. **(G)** Protein normalized peak areas of labeled SAM from ^13^C_5_-methionine (200µM, last 1h) tracing of cells treated as in (F). **(H)** Fractional enrichment of SAM isotopologues from^13^C_5_-methionine tracing as in (G). **(I)** Protein normalized peak areas of labeled cystathionine from^13^C_5_-methionine tracing as in (G). Data are plotted as mean ± SD for tracing studies and are representative of two independent experiments performed in triplicate. *p<0.05, **p<0.01, ***p<0.001, ****p<0.0001 (two-way ANOVA).

The methionine cycle produces SAM through the activity of the methionine adenosyltransferase (MAT) enzymes (MAT1 and MAT3 in liver [46]), and SAM is the methyl donor for a variety of cellular processes including histone, DNA, and protein methylation and phosphatidylcholine synthesis [47]. The methionine cycle is tightly coupled to both folate metabolism and the transsulfuration pathway (**Figure 5C**). Given the observed alterations in metabolites of the methionine cycle and transsulfuration pathway, we broadly assessed the expression of genes encoding enzymes within this metabolic network in response to insulin and *Atf4* knockdown. Genes of the cytosolic THF and methionine cycles were insensitive to insulin and ATF4 depletion in primary hepatocytes (**Figure 5D**). Notably, the MAT enzymes were not found to be transcriptionally regulated in this setting, in contrast to proliferating cells, where MAT2A has recently been found to be regulated by mTORC1 signaling via c-Myc [48]. Interestingly, only *Cth*, encoding an enzyme of the transsulfuration pathway, also known as cystathionine gamma lyase (CGL), was found to be significantly induced with insulin in an ATF4-dependent manner (**Figure 5D**). Like other targets of the mTORC1-ATF4 axis in primary hepatocytes (**Figure 1F**), the insulin-stimulated expression of *Cth* was also found to peak at 4h and be fully suppressed by rapamycin or Torin1 treatment (**Figure S4D**). The insulin- and mTORC1-mediated regulation of *Cth* expression was lost in primary hepatocytes derived from *LAtf4^KO^* mice, and *Cth* expression was restored to these cells with exogenous ATF4 (**Figure S4E,F**).

To determine if the insulin-mTORC1-ATF4 pathway influenced metabolic flux into the methionine cycle and transsulfuration pathways in primary hepatocytes, we utilized stable-isotope tracing. Indeed, insulin stimulated 3-^13^C_1_-serine tracing into labeled SAM (m+1) and cystathionine (m+1) in a manner that was attenuated by either mTOR inhibitors or siRNA knockdown of *Atf4* (**Figure 5E,F**). Of note, no change in the labeling of other metabolites of the methionine cycle, including methionine, S-adenosylhomocysteine (SAH), and homocysteine, were detected in this experiment (data not shown). To further assess the point of regulation of insulin- and ATF4-mediated SAM synthesis, we employed ^13^C_5_-methionine tracing to directly test involvement of methionine conversion to SAM via MAT isoforms. Robust SAM labeling (m+5) was detected (>70% fractional enrichment after 1 h labeling) but was unaffected in primary hepatocytes first stimulated with insulin or depleted of ATF4 (**Figure 5G,H**). Conversely, insulin stimulated an increase in labeling of the transsulfuration metabolites cystathionine (m+4) and α-ketobutyrate (m+4) from ^13^C_5_-methionine in a manner blunted by *Atf4* knockdown (**Figure 5I** and **Figure S4G**). Thus, insulin and ATF4 induce SAM synthesis through a mechanism independent of MAT regulation and stimulate the transsulfuration pathway downstream of the methionine cycle.

### 3.6. Insulin induces hepatocyte aspartate and alanine synthesis via ATF4

The steady state metabolomic profiling of primary hepatocytes with *Atf4* knockdown also showed that the non-essential amino acids aspartate and alanine were increased with insulin in an ATF4-dependent manner (**Figure 5A**). To determine whether this change reflects a stimulated increase in the synthesis of these amino acids, we employed stable isotope tracing with ^15^N-glutamine-amine labeling. Aspartate synthesis results from transamination of oxaloacetate by GOT1 in the cytosol and GOT2 in the mitochondria (**Figure S5A**). ^15^N-glutamine-amine flux into labeled aspartate (m+1) in primary hepatocytes was increased with insulin in a rapamycin-resistant, but Torin1-sensitive manner (**Figure S5B**), perhaps reflecting the more potent effects of Torin1 on ATF4 and its targets (**Figure 1**). One-hour treatment with cycloheximide increased the labeled pool of aspartate, indicating that a portion of newly synthesized aspartate is rapidly utilized for protein synthesis. Consistent with the steady state measurements (**Figure 5A,B**), stable-isotope tracing found that insulin stimulated an increase in aspartate synthesis in a manner ablated by siRNA knockdown of *Atf4* (**Figure S5C**). However, unlike other cell types [26, 49], the expression of neither *Got1* nor *Got2* were affected by ATF4 depletion in hepatocytes (**Figure S5D-E**). We next assessed alanine synthesis in primary hepatocytes, which results from transamination of pyruvate by GPT1 in the cytosol and GPT2 in the mitochondria (**Figure S5F**). Much like aspartate synthesis, ^15^N-glutamine-amine flux into labeled alanine (m+1) was increased with insulin in rapamycin-resistant, but Torin1-sensitive manner in primary hepatocytes (**Figure S6G**). Of note, cycloheximide treatment had minimal effects on the free pool of newly synthesized alanine. Also similar to aspartate, stable-isotope tracing found that insulin stimulated an increase in alanine synthesis in an ATF4-dependent manner (**Figure S6H**). Previous studies have linked ATF4 to the transcriptional regulation of *Gpt2* [26, 42, 49, 50], and *Atf4* knockdown decreased hepatocyte *Gpt2* expression under both basal and insulin-stimulated conditions (**Figure S6H**). However, unlike alanine synthesis, insulin did not stimulate a significant increase in *Gpt2* transcripts, suggesting that these two observations may be unrelated. Together, these results indicate that insulin-mTORC1 signaling stimulates aspartate and alanine synthesis in primary hepatocytes via ATF4, albeit through a currently unknown mechanism.

### 3.7. Feeding induces an ATF4-dependent transcriptional response in the mouse liver

Previous studies have challenged the *LAtf4^KO^* mice with stress stimuli that engage the ISR [28, 51], but there are no reports to date characterizing the transcriptional response to feeding in these mice. To first confirm that ATF4 was ablated in the livers of *LAtf4^KO^* mice, and to screen commercially available antibodies for specific recognition of ATF4 in the mouse liver, we treated *Atf4^fl/fl^* and *LAtf4^KO^* mice with the ER stress-inducing agent tunicamycin to stimulate a robust increase in hepatic ATF4 levels (**Figure S6A**). Indeed, tunicamycin failed to induce detectable ATF4 protein in the livers of *LAtf4^KO^* mice. It is also important to note that this analysis revealed that some widely used ATF4 antibodies recognize non-specific bands at the same molecular mass as ATF4 on immunoblots of mouse liver extracts. We next used the same daytime fasting and nighttime refeeding paradigm employed in Figure 1A,B. The *LAtf4^KO^* livers displayed comparable levels of mTORC1 activation to the *Atf4^fl/fl^* control livers upon refeeding, which correlated with a robust increase in both full length and processed forms of SREBP1c, a known effector of hepatic mTORC1 with feeding [4, 6, 20] (**Figure 6A**). It is worth noting that this result is counter to what was observed previously in livers from whole body *Atf4^-/-^* mice, which were reported to display a decrease in hepatic SREBP1c activation, measured by expression of its transcriptional targets [52, 53]. In addition to normal induction of mTORC1 signaling, the feeding-induced expression of the SREBP1c target *Fasn* and suppression of the gluconeogenic gene *Pepck* were similar between the *Atf4^fl/fl^* and *LAtf4^KO^* livers, indicative of a normal feeding response in the *LAtf4^KO^* livers (**Figure 6B**). Gene expression analysis confirmed our previous observation that feeding induces the expression of the serine synthesis pathway genes (*Phgdh*, *Psat1*, and *Psph*) and *Mtfhfd2* in the *Atf4^fl/fl^* livers, and this response was blunted in the *LAtf4^KO^* livers, providing genetic evidence of their regulation by feeding through ATF4 activation.

**Figure 6.**
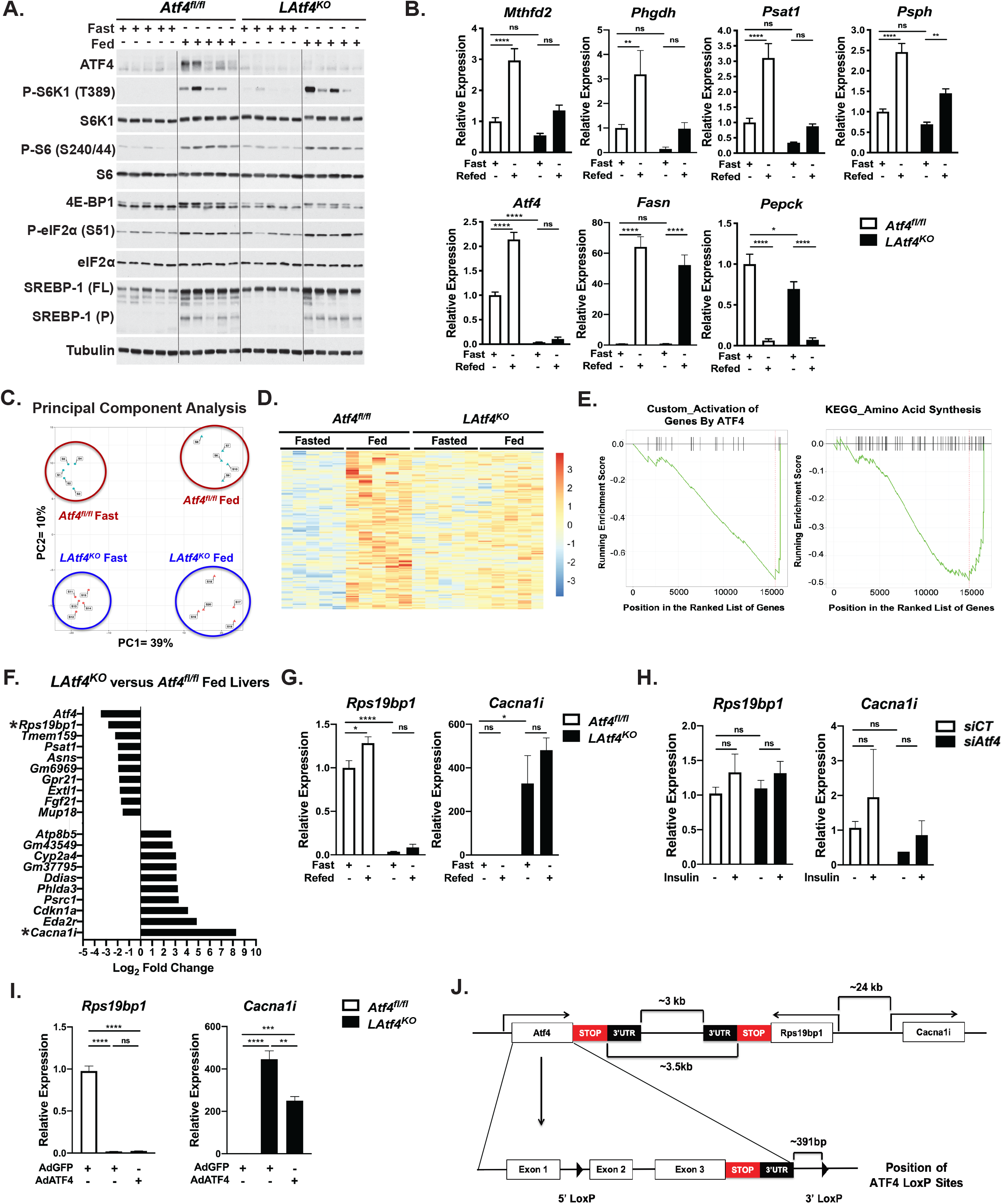
Characterization of feeding induced transcriptional response in *LAtf4^KO^* mice. **(A,B)** Eight-week old *Atf4^fl/fl^* and *LAtf4^KO^* male mice were fasted for 12h and refed a high carbohydrate diet for 6h (n=5/group). Immunoblots (A) and gene expression analysis (B) of *Atf4^fl/fl^* and *LAtf4^KO^* livers. Gene expression data are plotted as mean ± SEM relative to the *Atf4^fl/fl^* fasted group (n=5 mice/group). **(C)** Principal component analysis of normalized transcripts from RNA-seq of *Atf4^fl/fl^* and *LAtf4^KO^* livers treated as in (A). **(D)** Heat map of normalized transcripts from RNA-seq data significantly (p_adj_<0.05) induced with feeding in the *Atf4^fl/fl^*, but not in the *LAtf4^KO^* livers. **(E)** Gene set enrichment analysis of the differentially expressed transcripts in fed *LAtf4^KO^* versus *Atf4^fl/fl^* livers. **(F)** Top-10 most upregulated and down-regulated transcripts plotted as log_2_ fold change in *LAtf4^KO^* versus *Atf4^fl/fl^* livers from fed mice. Asterisks indicate top hits. **(G)** Gene expression analysis of *Atf4^fl/fl^* and *LAtf4^KO^* livers treated and plotted as in (A). **(H)** Gene expression analysis of primary mouse hepatocytes transfected with control (siCT) or *Atf4*-targetting siRNAs followed by overnight serum starvation and treatment with 100nM insulin for 8h. Data are plotted as mean ± SEM relative to the *siCT*-transfected cells (n=3 independent experiments). **(I)** Gene expression analysis of primary mouse hepatocytes from *Atf4^fl/fl^* and *LAtf4^KO^* mice infected with AdGFP or AdATF4 (MOI=10) for 24h. Data from a representative experiment are plotted as mean± SEM relative to the *Atf4^fl/fl^* AdGFP cells (n=3). **(J)** Schematic of the *Atf4* locus and proximal genes on mouse chromosome 15, along with the modified *Atf4^fl/fl^* locus. *p<0.05, **p<0.01, ***p<0.001, ****p<0.0001 (two-way ANOVA (B, G, I) or one-way ANOVA (H)).

Given that whole body *Atf4^-/-^* mice display growth defects and lower body weights [54], we assessed physiological parameters in the *LAtf4^KO^* mice. We found that body weights of the *Atf4^fl/fl^* and *LAtf4^KO^* mice were similar at 8 weeks, however there was a small, but statistically significant, body weight reduction in the *LAtf4^KO^* mice at 26 weeks of age (**Figure S6B**). Furthermore, there were no significant differences observed in systemic glucose and insulin tolerance between *Atf4^fl/fl^* and *LAtf4^KO^* mice at 26 weeks of age on a chow diet (**Figure S6C**), consistent with previous observations [51].

To more broadly define the transcriptional output of ATF4 downstream of physiological mTORC1 activation, we performed RNA-seq analysis on *Atf4^fl/fl^* and *LAtf4^KO^* livers with fasting and refeeding. Principal component analysis of the normalized, differentially expressed transcripts revealed that transcriptional changes with fasting and feeding produce the most robust changes between groups, accounting for approximately 39% of the overall variance, whereas genotype accounted for approximately 10% of the variance (**Figure 6C**). We focused on the subset of genes that were significantly induced with feeding in the *Atf4^fl/fl^* livers but not in the *LAtf4^KO^* livers (141 genes, log_2_FC>1, p<0.05; heatmap in **Figure 6D** and **Supplementary Table 5**). Gene set enrichment analysis revealed that previously defined ATF4 target genes involved in amino acid synthesis were enriched in the genes most downregulated in the *LAtf4^KO^* livers relative to the *Atf4^fl/fl^* livers with feeding (**Figure 6E**). These data indicate that the recently recognized role of ATF4 in amino acid metabolism downstream of mTORC1 signaling obtained from cell culture models is active in the liver in response to feeding [25, 26].

Notably, gene set enrichment analysis revealed that p53 target genes are enriched in the list of upregulated genes in *LAtf4^KO^* livers relative to the *Atf4^fl/fl^* livers with feeding (**Figure S7A**). In line with this observation, *LAtf4^KO^* livers from both fasted and fed mice displayed a robust increase p53 protein levels relative to the *Atf4^fl/fl^* livers (**Figure S7B**). Moreover, the p53 target genes *Cdkn1a* (p21), *Ddias*, *Phlda3*, *Psrc1*, and *Eda2r* were all confirmed by qRT-PCR to be significantly upregulated in the *LAtf4^KO^* livers relative to the *Atf4^fl/fl^* livers (**Figure S7C**). To determine if this activation of p53 was intrinsic to hepatocytes, we isolated primary hepatocytes from the livers of *Atf4^fl/fl^* and *LAtf4^KO^* mice. Indeed, primary hepatocytes from *LAtf4^KO^* mice also displayed upregulation of representative p53 target genes and p53 protein levels (**Figure S7D,E**). However, exogenous expression of ATF4 failed to suppress p53 or its gene targets in the *LAtf4^KO^* hepatocytes, while predictably restoring expression of the canonical ATF4 target *Asns* (Figure S7D-E). Furthermore, the expression of p53 target genes was unaffected by siRNA-mediated knockdown of ATF4 in primary hepatocytes (**Figure S7F**). Thus, the strong induction of p53 and its transcriptional targets upon loss of liver ATF4 may not be an immediate or direct effect of ATF4 loss in hepatocytes.

### 3.8. Secondary effects of Cre-mediated recombination at the *Atf4^fl/fl^* locus

Examination of the top ten most downregulated genes in the *LAtf4^KO^* livers revealed that, along with established ATF4 targets (*Psat*, *Asns*, *Fgf21*), *Rps19bp1*, a previously described ribosomal protein S19-binding protein linked, among other things, to the regulation of protein synthesis [55], was second only to *Atf4* among most downregulated genes (**Figure 6F**). Of note, siRNA-mediated knockdown of Rps19bp1 in primary hepatocytes did not alter the expression of p53 target genes (**Figure S7F**; discussed below). Five of the top ten most upregulated genes in the *LAtf4^KO^* livers relative to *Atf4^fl/fl^* controls were established targets of p53, but the most increased RNA-seq transcript reads were from the *Cacna1i* gene, encoding a brain-specific calcium transporter (Cav3.3) previously linked to schizophrenia [56]. Validation experiments demonstrated that *Rps19bp1* expression was completely abrogated in the *LAtf4^KO^* livers, while *Cacna1i* is upregulated ∼400-500 fold in the *LAtf4^KO^* livers relative to the control livers (**Figure 6G**). However, acute silencing of *Atf4* with siRNAs in primary hepatocytes did not significantly alter *Rps19bp1* or *Cacna1i* expression, suggesting that the effects are specific to the *LAtf4^KO^* context (**Figure 6H**). Indeed, the gene expression findings from the liver were recapitulated in primary hepatocytes cultured from these mice, but exogenous re-expression of ATF4 in *LAtf4^KO^* hepatocytes failed to rescue the lost expression of *Rps19bp1*, while modestly but significantly reducing *Cacna1i* expression (**Figure 6I**). Examination of the *Atf4* locus on mouse chromosome 15 revealed that both *Rps19bp1* and *Cacna1i* were proximal to *Atf4*. The end of the *Atf4* 3’UTR-encoding region is approximately 3 kb from that of the convergently transcribed *Rps19bp1* gene, an intergenic region where the 3’ LoxP site is located in the *Atf4^fl/fl^* allele (**Figure 6J**). In addition, the start codon of the *Cacna1i* gene lies approximately 24 kb from that of the *Rps19bp1* gene. It is worth noting that expression of genes in loci upstream of *Atf4*, such as *Mgat3*, were not perturbed by *Cre*-mediated recombination in the *LAtf4^KO^* livers (data not shown). Thus, the extreme alterations to *Rps19bp1* and *Cacna1i* expression upon Cre-mediated deletion of exons 2 and 3 of *Atf4* are likely a secondary result of disruptions to the chromosome 15 architecture in this region (discussed further below).

## 4. Discussion

Recent studies demonstrate that ATF4 is a downstream target of mTORC1 signaling in proliferating cells [24–26]. In non-proliferative settings such as the liver, ATF4 function has been largely studied in the context of stress, including amino acid limitation, ER stress, and in models of obesity and fatty liver [28, 51, 57–59]. Here, we define ATF4 as a novel metabolic effector of physiological mTORC1 activation with feeding in the liver and insulin in primary hepatocytes. While the mTORC1-ATF4 axis has recently been found to be activated in the pancreatic islets of mice with β cell-specific genetic ablation of the secretory peptidase Furin [60], our findings place ATF4 as a downstream target of mTORC1 in a non-proliferative metabolic tissue activated in response to hormonal cues. Dynamic functional regulation of hepatic mTOR signaling with fasting and feeding was reported in neonatal pigs two decades ago, with protein synthesis as the primary metabolic output [8]. However, to date, only SREBP1c has been implicated as a downstream transcriptional effector of mTORC1 signaling in the liver activated in response to feeding and insulin [4, 6, 20], which pales in comparison to the extensive transcriptional network described for mTORC1 signaling in proliferating cells [61]. While more work is necessary to fully define how mTORC1 alters hepatic metabolism in response to feeding, this current study demonstrates that the canonical stress-responsive transcription factor ATF4 can be alternatively activated downstream of mTORC1 the liver to control specific metabolic processes, including amino acid metabolism.

Given the robust mTORC1-ATF4-mediated regulation of genes of the serine/glycine synthesis and one-carbon metabolism pathways, which provide essential substrates for *de novo* purine synthesis, observed in the intact liver and isolated hepatocytes in this study, we hypothesized that this signaling axis would control these processes in response to insulin. Compared to other tissues, the liver has a massive protein synthetic capacity, as approximately 85-90% of the protein content of blood serum originates in this organ [62]. Such a demand for protein synthesis would necessitate a robust program of ribosome biogenesis that includes the synthesis of nucleotides essential for rRNA production. Indeed, we observed that mTORC1 promotes the *de novo* synthesis of both purine and pyrimidine nucleotides in primary hepatocytes, consistent with previous studies in proliferating cells [11, 12, 24]. However, insulin and mTORC1 stimulated these processes in an ATF4-independent manner. A previous study in cell culture models identified other transcription factors, in addition to ATF4, that stimulated *de novo* purine synthesis in response to mTORC1 activation, which could underlie this regulation in hepatocytes [63]. We provide evidence that both feeding and insulin stimulate hepatic CAD phosphorylation in an mTORC1-dependent manner, an activating modification on this enzyme leading to increased flux through the *de novo* pyrimidine synthesis pathway [11, 12]. Further studies are necessary to better elucidate the mechanism(s) of mTORC1-driven *de novo* nucleotide synthesis in the liver, but the ability of insulin to stimulate this process occurs independent of ATF4, at least in primary hepatocytes. Lastly, it is worth noting that the fractional enrichment of *de novo* synthesized nucleotides in total hepatocyte RNA measured in our study was much lower than expected. Thus, factors influencing ribosome synthesis and turnover in the liver, along with specific effects on other RNA species such as tRNAs, which are also critical to support protein synthesis, are important areas for future investigation.

ATF4 is known to induce the expression of genes encoding nearly every enzyme of non-essential amino acid synthesis, a function documented downstream of both the ISR and mTORC1 activation in other settings [22, 24–26, 42]. Despite evident regulation of the serine synthesis genes PHGDH, PSAT1, and PSPH by insulin signaling through mTORC1 and ATF4 in primary hepatocytes, we failed to detect any regulated flux through the serine synthesis pathway, which was extremely low under all conditions tested. Our findings are consistent with data from a recent study that showed the fractional enrichment of serine (m+3) synthesized from ^13^C_6_-glucose in the liver was minimal (<1%) [64]. Thus, despite clear regulation of the serine synthesis enzymes, hepatocytes might rely predominantly on exogenous serine. In contrast to serine synthesis, we did observe insulin and ATF4-dependent synthesis of aspartate and alanine, but the genes encoding the aminotransferase enzymes for aspartate and alanine synthesis, which have been established as transcriptional targets of ATF4 in other settings [25, 26, 42, 49] were not regulated by insulin or ATF4 in hepatocytes. These data highlight the importance of measuring metabolic flux, as changes in enzyme expression do not always reflect changes in metabolic activity, which is also strongly influenced by allosteric regulation and concentrations of metabolic substrates, cofactors, and products. Future studies are warranted to determine the mechanism by which ATF4 influences aspartate and alanine synthesis in response to insulin and mTORC1 signaling in primary hepatocytes.

Unbiased metabolomics and subsequent metabolic flux analyses demonstrated that the hepatocyte insulin-mTORC1-ATF4 pathway stimulates SAM production within the methionine cycle, as well as the transsulfuration pathway, which shunts off of the methionine cycle. While a recent study demonstrated that mTORC1 regulates SAM synthesis in proliferating cells through c-Myc-dependent control of *Mat2a* expression [48], the levels of transcripts encoding the MAT enzymes were unaffected by insulin signaling in hepatocytes. Hepatic MAT activity can also be attenuated by reactive oxygen species [46], which are elevated in other models of ATF4 deficiency [22]. However, methionine tracing into SAM, as a more direct assay of MAT activity, indicated that this enzymatic step is unlikely to be the major point of regulation by insulin-ATF4 signaling in hepatocytes. Importantly, the methionine cycle is calibrated to the THF cycle via SAM allosterically inhibiting MTHFR, which serves as the entry point of serine-derived one carbon units into the cycle [47]. It is possible that the defect in insulin-stimulated SAM synthesis observed with ATF4 knockdown in hepatocytes stems from a perturbation in redox that affects MTHFR activity or from other defects in the THF cycle. We also observed a striking insulin-stimulated increase in the synthesis of the transsulfuration pathway intermediate cystathionine downstream of mTORC1 and ATF4. Within the transsulfuration pathway, insulin, mTORC1, and ATF4 were found to regulate expression of the second enzyme in the pathway, cystathionine γ-lyase, encoded by *Cth*, but not the first enzyme off of the methionine cycle, cystathionine β-synthase, encoded by *Cbs*, consistent with studies on ATF4 activated as part of the ISR [65]. Interestingly, CBS activity can be allosterically activated by SAM [66], raising the possibility that the SAM and cystathionine changes might be mechanistically linked. However, insulin signaling leading to ATF4 activation did not influence methionine tracing into SAM but did stimulate methionine tracing into cystathionine, indicating that the effects of ATF4 on these two metabolites are separable in hepatocytes. Given that the insulin-mTORC1-ATF4 pathway induces *Cth* expression in hepatocytes, it seems likely that the observed changes in steady state levels and metabolic flux into cystathionine are mediated through this regulation. This is also consistent with the fact that α-ketobutyrate produced in the CTH reaction is similarly regulated. It should be noted that technical issues prevented us from reliably measuring the synthesis of cysteine, the other product of this reaction. As *Cth* was the only gene in these interconnected metabolic pathways found to be induced with insulin in an ATF4-dependent manner, it is possible that regulation of the transsulfuration pathway at the CTH step influences both the THF and methionine cycles to account for increased methionine disposal through this route. This regulation of the transsulfuration pathway has potentially important metabolic consequences in the liver, as its product cysteine is required for the synthesis of protein, glutathione, and taurine, the latter of which contributes to hepatic bile acid production [66, 67].

While we were able to identify and validate feeding-induced ATF4 target genes in the liver using the *LAtf4^KO^* model, a limitation of our study is the dramatic alterations to *Rps19bp1* and *Cacna1i* expression found to accompany Cre-mediated deletion of *Atf4* in this model. These changes make it more difficult to definitively assign specific phenotypes to loss of ATF4 function in the liver of these mice. For example, while a p53-mediated stress response has been observed in other settings of ATF4 loss [68–70], it is possible that the robust upregulation of p53 and its gene targets observed in the *LAtf4^KO^* livers and cultured hepatocytes could stem from loss of *Rps19bp1* expression. The protein encoded by this gene, also known as active regulator of SIRT1 (AROS), acts with SIRT1 to deacetylate and inactivate p53 [71]. However, transient knockdown of *Rps19bp1/Aros* was not sufficient to induce the expression of p53 target genes in primary hepatocytes in our study. Importantly, similar alterations to the expression of *Rps19bp1* and *Cacna1i* have recently been observed in RNA-Seq studies of skeletal muscle from muscle-specific *Atf4* knockout mice (*mAtf4^KO^*) (C.M. Adams, *personal communication*). It is worth noting that *mAtf4^KO^* muscle does not exhibit increased expression of p53 or its gene targets [72, 73], indicating that p53 activation does not coincide with reduced *Rps19bp1* expression in all settings. The apparent increase in *Cacna1i* transcripts in *mAtf4^KO^* muscle has been characterized to result from a non-natural fusion transcript consisting of *Atf4* exon 1 fused to *Cacna1i* exon 2, which produces an out-of-frame fusion transcript, without alterations to the normal full-length *Cacna1i* mRNA. Thus, it is unlikely that the effects on this transcript yield functional consequences. Nonetheless, future studies employing these and other mouse models that genetically ablate *Atf4* should consider the potential for similar effects on neighboring genes.

In summary, we describe ATF4 as a novel metabolic effector of hepatic mTORC1 signaling in response to insulin. While there are likely other downstream mediators of the mTORC1-dependent feeding response yet to be defined, this study advances our understanding of how mTORC1 exerts metabolic control in a physiological setting. Notably, hepatic mTORC1 and ATF4 are both upregulated in obesity [74–76]. It will be interesting in future studies to determine the relative contribution of stress signaling and mTORC1 signaling to ATF4 activation in this setting, perhaps revealing how adaptive regulation of ATF4 with feeding becomes maladaptive when chronically engaged in disease.

## Acknowledgements

We thank Tracy G. Anthony for advice on ATF4 antibodies and mouse models and Tiffany Horng, Sudha Biddinger, Nada Kalaany, Gyan Prakash, Matthew Miller, and members of the Manning lab for advice and technical assistance. This research was supported by grants from the NIH: Joslin Diabetes Center T32-NK007260 (V.B.), R35-CA197459 (B.D.M.), P01-CA120964 (B.D.M. and J.A.), and R01-AR071762 and R01-AG060637 (C.M.A.); the Congressionally Directed Medical Research Program on Tuberous Sclerosis Complex award no. W81XWH-18-1-0659 (B.D.M.); U.S. Department of Veteran Affairs grant I01BX00976 (C.M.A.); and a research grant from Zafgen (B.D.M.). These funders were not involved in the design, execution, or interpretation of this study. C.M.A. is a shareholder, director, and officer of Emmyon, Inc. B.D.M. is a shareholder and scientific advisory board member of Navitor Pharmaceuticals. All other authors declare no competing financial interests.

## Author Contributions

V.B. conceived of the project, performed experiments and data analysis and wrote the manuscript. Y.C., K.K., G.H. and J.H.H performed experiments. V.B-B. and S.H.S. analyzed the RNA-sequencing. J.M.A. and I.B.S. performed mass spectrometry. C.M.A. supplied the *Atf4^fl/fl^* mice and critical insights into ATF4. B.D.M. conceived of and supervised the project and wrote the manuscript.

**Supplementary Table 1.** List of commercially available ATF4 antibodies tested in *in vitro* and *in vivo*.

**Supplementary Table 2.** Selected reaction monitoring (SRM) table for stable isotope tracing experiments.

**Supplementary Table 3.** Primer sequences used for qPCR analysis.

**Supplementary Table 4.** Peak areas and p-values for metabolites in Figure 5A heatmap. Data sorted by fold change of siCT +Insulin/siCT -Insulin. P-value determined by Student’s t-test.

**Supplementary Table 5.** List of genes induced with feeding in the *Atf4^fl/fl^* (log_2_FC>1, p_adj_<0.05), but not the *LAtf4^KO^* mouse livers.

---

**Supplementary Figure 1.**
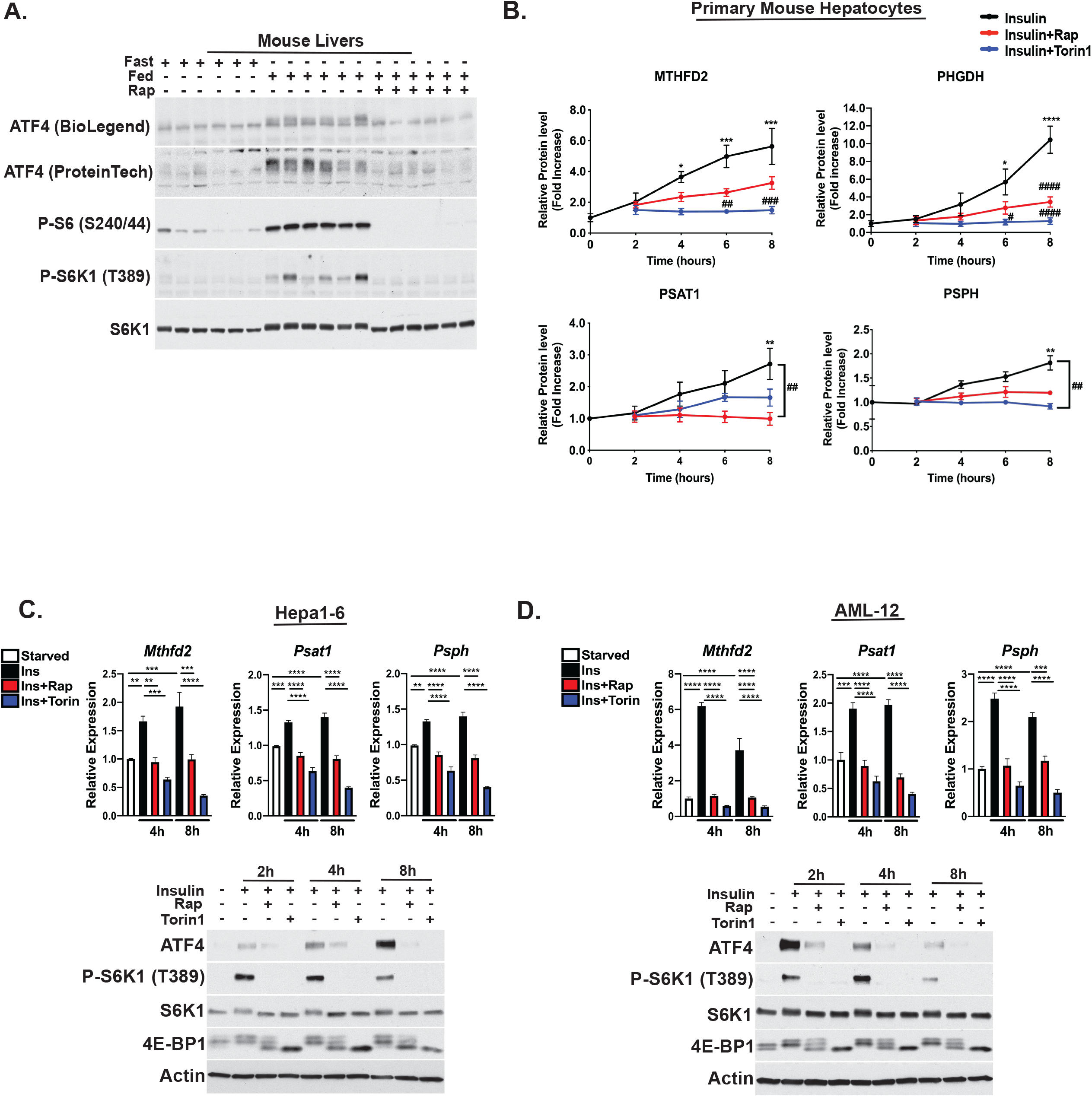
Feeding and insulin induce hepatic ATF4 and established gene targets via mTORC1 signaling. **(A)** Eight-week old male mice were fasted for 12h and refed a high carbohydrate diet for 12h following pretreatment with vehicle or 10mg/kg rapamycin (n=6/group). Liver lysates were immunoblotted to assess mTORC1 signaling and ATF4 protein levels. (**B)** LICOR quantification of immunoblots from primary mouse hepatocytes serum starved overnight and subsequently treated with 100nM insulin following a 30-minute pretreatment with vehicle (DMSO), 20nM rapamycin, or 750nM Torin1 for the indicated time points. Data are plotted as ± SEM relative to the serum starved samples (n=4 independent experiments). **(C,D)** Hepa1-6 and AML-12 cells were serum starved overnight and treated with 100nM insulin following a 30-minute pretreatment with vehicle (DMSO), 20nM rapamycin, or 250nM Torin1 for the indicated time points. *p<0.05, **p<0.01, ***p<0.001, ****p<0.0001. In panel C, * denote comparisons to serum starved samples at each insulin time point and ^#^p<0.05, ^##^p<0.01, ^###^p<0.001, ^####^p<0.0001 for comparison of insulin versus rapamycin or Torin1 at each time point (one-way ANOVA).

**Supplementary Figure 2.**
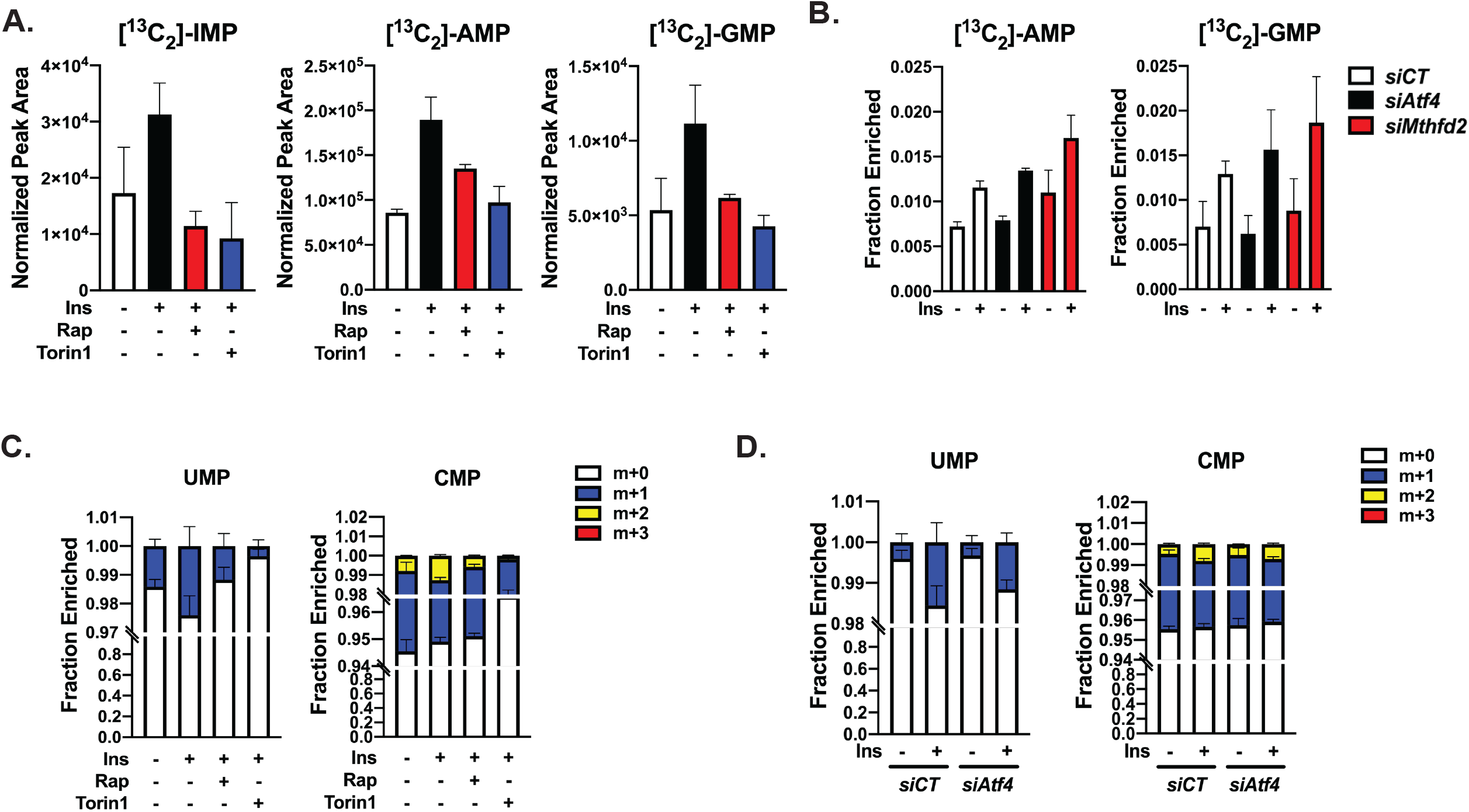
mTORC1 controls *de novo* nucleotide synthesis in an ATF4-independent manner. **(A)** Protein normalized peak areas of 3-^13^C-serine tracing (400µM, last 1h) into labeled purine nucleotide pools in primary mouse hepatocytes serum starved overnight and subsequently treated with 100nM insulin following a 30-minute pretreatment with vehicle (DMSO), 20nM rapamycin, or 500nM Torin1 for 8h. Data are plotted as mean ± SD and are representative of two independent experiments performed in triplicate. **(B)** Protein normalized peak areas of 3-^13^C-serine tracing (400µM, last 1h) into labeled purine nucleotide pools in primary mouse hepatocytes transfected with control (siCT) siRNAs or those targeting *Atf4* or *Mthfd2* followed by overnight serum starvation and treatment with 100nM insulin for 8h. Data are plotted as mean ± SD and are representative of two independent experiments performed in triplicate. **(C)** ^15^N-glutamine-amide tracing into RNA-derived pyrimidine nucleotide isotopologues in primary mouse hepatocytes serum starved overnight and subsequently treated with 10nM insulin following a 30 minute pretreatment with vehicle (DMSO), 20nM rapamycin, or 500nM Torin1 for 8h in medium containing 2mM ^15^N-glutamine-amide tracer. Fractional enrichment is plotted as mean ± SD and is representative of two independent experiments performed in triplicate. **(D)** ^15^N-glutamine-amide tracing into RNA-derived purine nucleotide isotopologues in primary mouse hepatocytes transfected with control (siCT) or *Atf4*-targetting siRNAs followed by overnight serum and treatment with 10nM insulin medium containing 2mM ^15^N-glutamine-amide tracer for 8h. Fractional enrichment is plotted as mean ± SD and is representative of three independent experiments performed in triplicate.

**Supplementary Figure 3.**
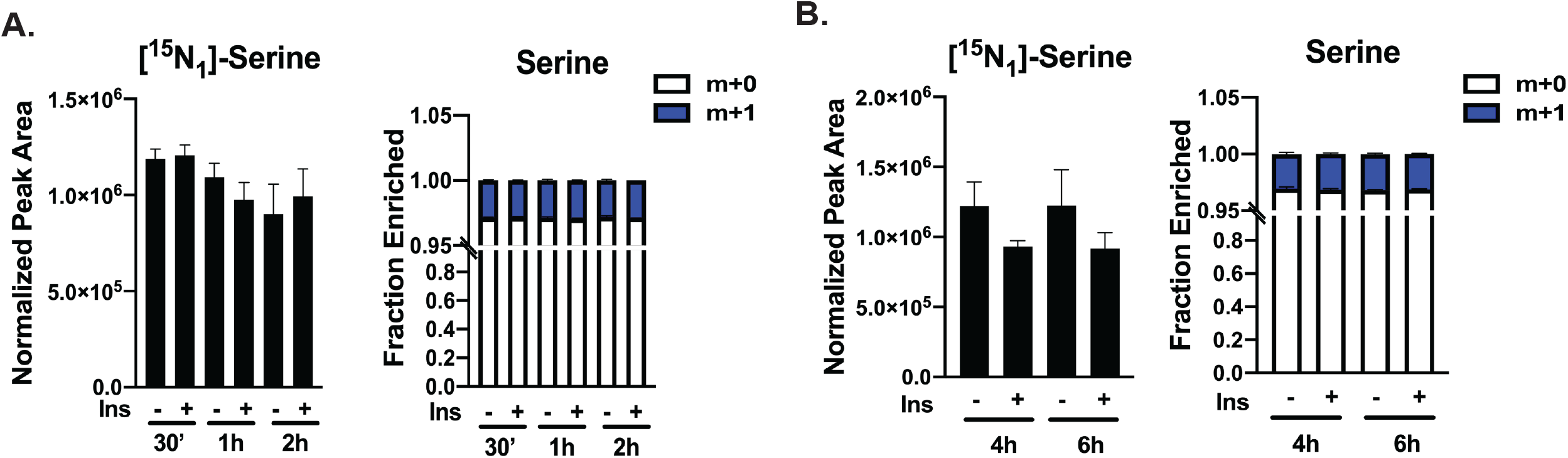
Effects of labeling time on measurements of serine synthesis in primary hepatocytes. **(A-B)** Protein normalized peak areas of ^15^N-glutamine-amine tracing into serine in primary mouse hepatocytes serum starved overnight and subsequently treated with 100nM insulin for 8h. Tracing with 2mM ^15^N-glutamine-amine was performed for indicated time points. Peak areas and fractional enrichment of serine isotopologues are plotted as mean ± SD.

**Supplementary Figure 4.**
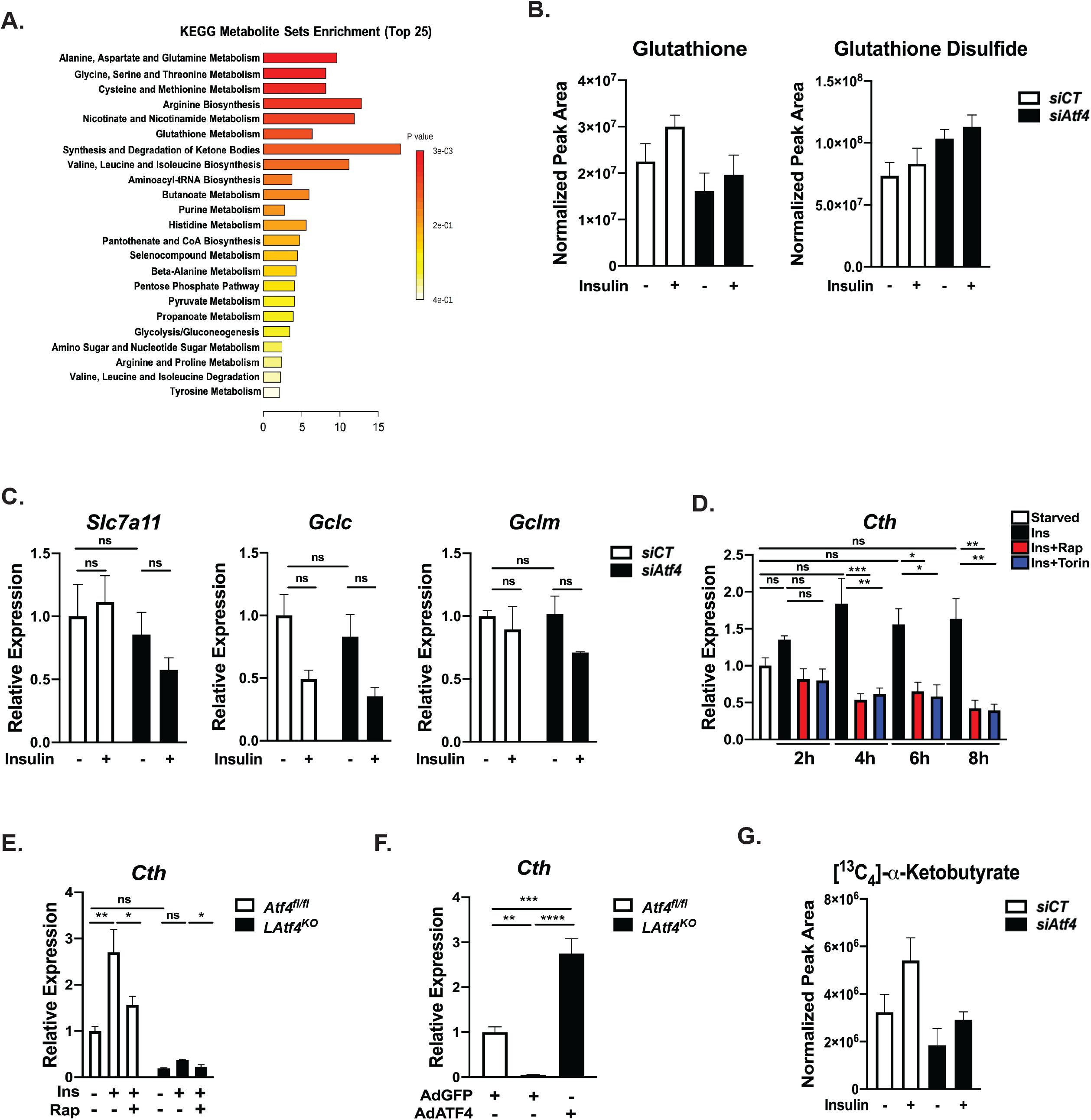
Insulin-mTORC1-ATF4 signaling regulates *Cth* and the transsulfuration pathway in primary hepatocytes. **(A)** Enrichment analysis of metabolites from Figure 4A, showing the enrichment score for the top 25 KEGG metabolite sets. **(B)** Steady state levels of glutathione and glutathione disulfide from primary mouse hepatocytes transfected with control (siCT) or *Atf4*-targetting siRNAs followed by overnight serum starvation and treatment with 100nM of insulin for 6h. **(C)** Gene expression analysis in primary mouse hepatocytes transfected and in (B) followed by overnight serum starvation and treatment with 100nM insulin for 8h. Data are plotted as mean ± SEM relative to *siCT* serum-starved cells (n=3 independent experiments). **(D)** Gene expression analysis of primary mouse hepatocytes serum starved overnight and subsequently treated with 100nM insulin following a 30-minute pretreatment with vehicle (DMSO), 20nM rapamycin, or 750nM Torin1 for the indicated time points. Data are plotted as mean ± SEM relative to serum-starved cells (n=4 independent experiments). **(E)** Gene expression analysis of *Atf4^fl/fl^* and *LAtf4^KO^* primary mouse hepatocytes serum starved overnight and subsequently treated with 100nM insulin for 6h following a 30-minute pretreatment with vehicle (DMSO) or 20nM rapamycin. Data from a representative experiment are plotted as mean± SEM (n=3) relative to serum-starved *Atf4^fl/fl^* cells. **(F)** Gene expression analysis of *Atf4^fl/fl^* and *LAtf4^KO^* primary mouse hepatocytes infected with AdGFP or AdATF4 (MOI=10) for 24h. Data from a representative experiment are plotted as mean± SEM (n=3) relative to *Atf4^fl/fl^* AdGFP cells. **(G)** Protein normalized peak areas of labeled alpha-ketobutyrate (m+4) from the ^13^C_5_-methionine tracing experiment in Figure 5. Data are plotted as mean± SD performed in triplicate. *p<0.05, **p<0.01, ***p<0.001, ****p<0.0001 (two-way ANOVA (C, E) or one-way ANOVA (D,F)).

**Supplementary Figure 5.**
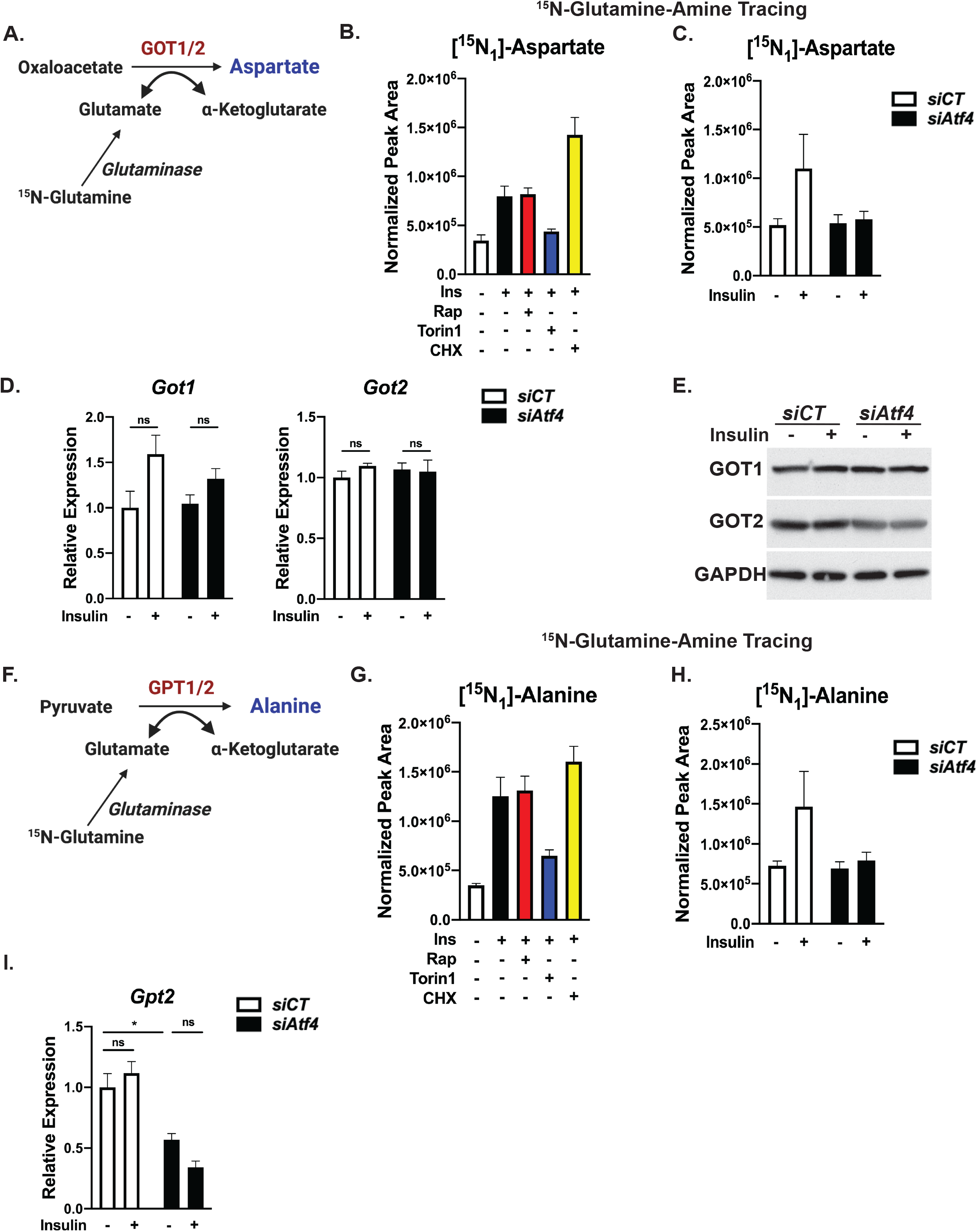
Insulin and ATF4 induce aspartate and alanine synthesis in primary hepatocytes. (**A**) Schematic of the glutamic-oxaloacetic transaminase (GOT) reaction to synthesize aspartate. (**B**) ^15^N-glutamine-amine tracing (2mM, last 1h) into labeled aspartate (m+1) in primary mouse hepatocytes serum starved overnight and subsequently treated with 100nM insulin following a 30-minute pretreatment with vehicle (DMSO), 20nM rapamycin, or 750nM Torin1 for 8h, or cycloheximide (50µM) for the last 1h. Data are plotted as mean ± SD and are representative of two independent experiments performed in quadruplicate. **(C)** Protein normalized peak areas of ^15^N-glutamine-amine tracing (2mM, last 1h) into labeled aspartate (m+1) in primary mouse hepatocytes transfected with control (siCT) or *Atf4*-targetting siRNAs followed by overnight serum starvation then treatment with 100nM insulin. Data are plotted as mean ± SD and are representative of two independent experiments performed in quadruplicate. **(D)** Gene expression analysis in primary mouse hepatocytes transfected with control (siCT) or *Atf4*-targetting siRNAs followed by overnight serum starvation and treatment with 100nM insulin for 8h. Data are plotted as mean ± SEM relative to serum-starved *siCT* cells (n=3 independent experiments). **(E)** Immunoblot analysis of primary mouse hepatocytes treated as in (D). **(F)** Schematic of the glutamic-pyruvic transamination (GPT) reaction to synthesize alanine. **(G)** Protein normalized peak areas of ^15^N-glutamine-amine tracing (2mM, last 1h) into labeled alanine (m+1) in primary mouse hepatocytes treated, and with data plotted, as in (B). **(H)** Protein normalized peak areas of ^15^N-glutamine-amine tracing into labeled alanine (m+1) in primary mouse hepatocytes treated, and with data plotted, as in (C). Gene expression analysis in primary mouse hepatocytes treated, and with data plotted, as in (D). *p<0.05, **p<0.01, ***p<0.001, ****p<0.0001 (two-way ANOVA).

**Supplementary Figure 6.**
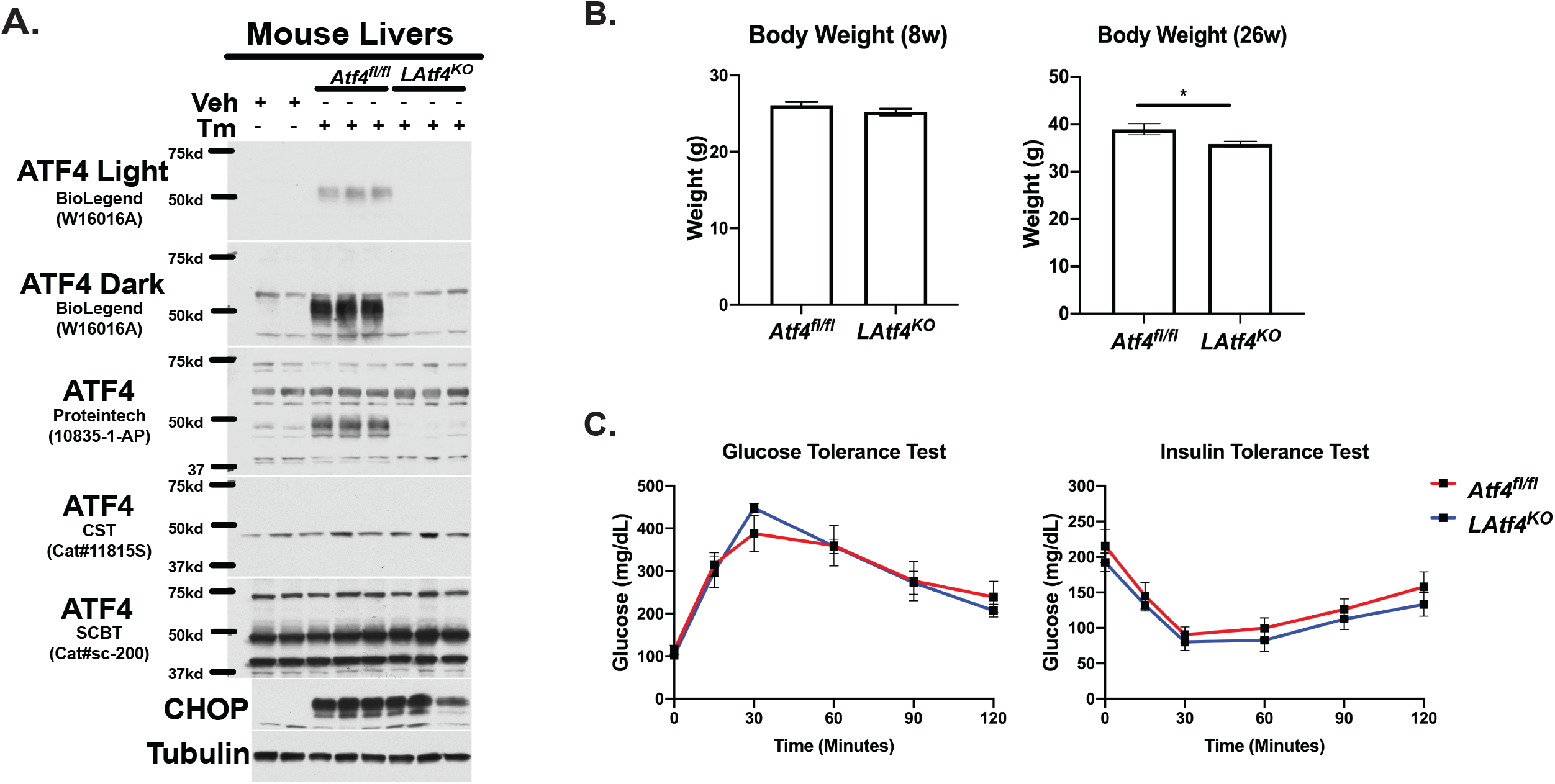
Physiological characterization of *LAtf4^KO^* mice. **(A)** Immunoblot analysis of livers from *Atf4^fl/fl^* and *LAtf4^KO^* mice injected with vehicle or 1mg/kg tunicamycin for 6h to test specificity of the indicated commercially available ATF4 antibodies. **(B)** Mean ± SEM body weights of *Atf4^fl/fl^* and *LAtf4^KO^* male mice at 8 weeks (n=15 mice/group) and 26 weeks of age (n=6 *Atf4^fl/fl^*, n=7 *LAtf4^KO^*) are plotted. **(C)** Glucose and insulin tolerance tests in male *Atf4^fl/fl^* (n=6) and *LAtf4^KO^* (n=7) mice at 26 weeks of age. *p<0.05, Students t-test (B).

**Supplementary Figure 7.**
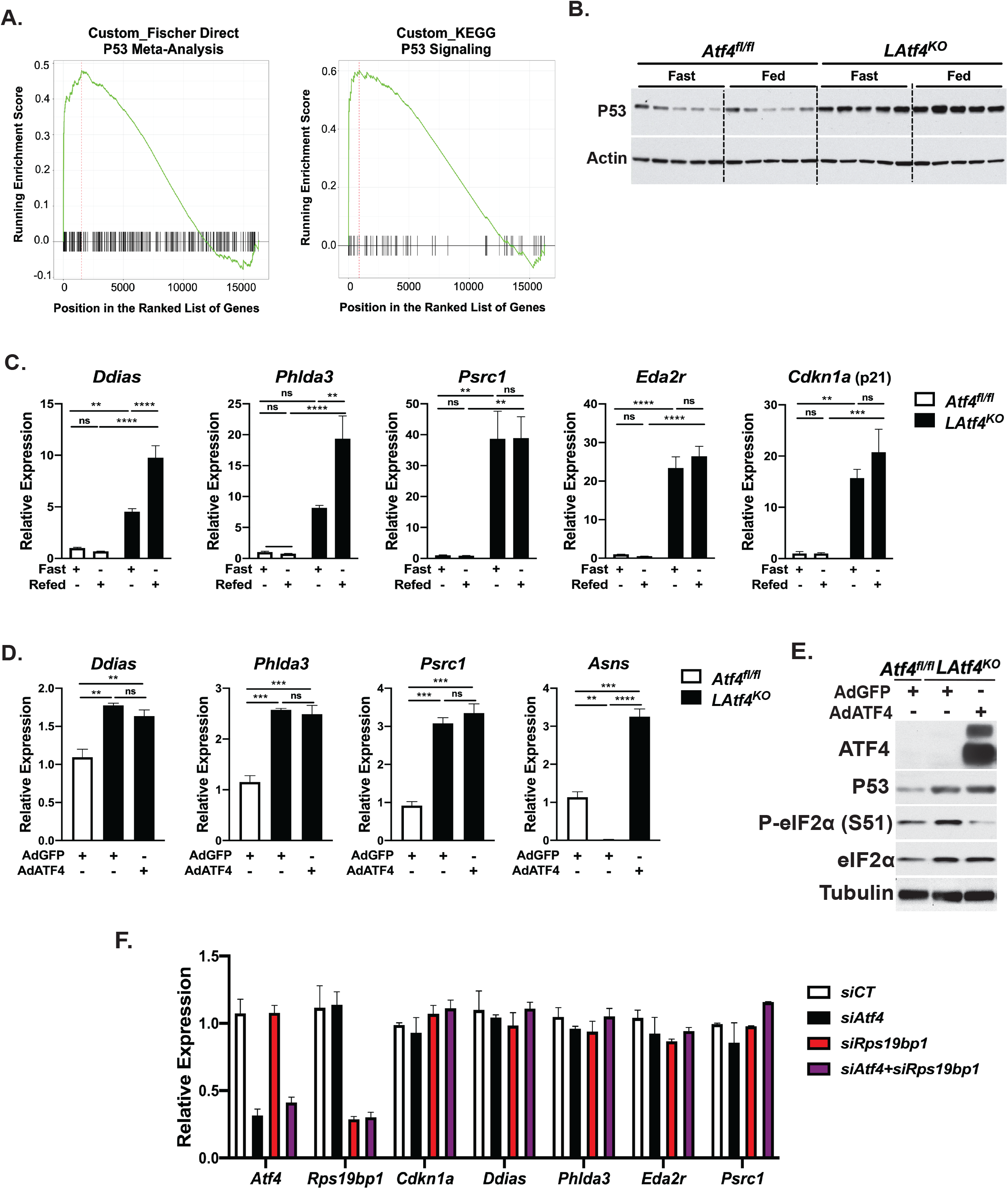
Upregulation of p53 and p53-dependent gene expression in *LAtf4^KO^* livers and primary hepatocytes. **(A)** Gene set enrichment analysis of the differentially expressed transcripts in fed *LAtf4^KO^* versus *Atf4^fl/fl^* livers. **(B)** Immunoblot of livers from eight-week old *Atf4^fl/fl^* and *LAtf4^KO^* mice subjected to a 12h fast followed by 6h refeeding (n=5 mice/group). **(C)** Gene expression analysis in livers from the mice in (E). Data are plotted as mean ± SEM relative to the *Atf4^fl/fl^* fasted group (n=5 mice/group). **(D)** Gene expression analysis of primary mouse hepatocytes from *Atf4^fl/fl^* and *LAtf4^KO^* mice infected with AdGFP or AdATF4 (MOI=10) for 24h. Data from a representative experiment are plotted as mean ± SEM relative to *Atf4^fl/fl^* AdGFP cells (n=3). **(E)** Immunoblot analysis of cells treated as in (G). **(F)** Gene expression analysis in primary mouse hepatocytes transfected with control (siCT), *Atf4*, or *Rps19bp1*-targeting siRNAs for 48h. Data are plotted as mean ± SEM relative to *siCT* cells (data are representative of two independent experiments). *p<0.05, **p<0.01, ***p<0.001, ****p<0.0001 (two-way ANOVA (F) or one-way ANOVA (G)).

